# Differences in maternal diet fiber content influence patterns of gene expression and chromatin accessibility in fetuses and piglets

**DOI:** 10.1101/2024.08.13.607725

**Authors:** Smahane Chalabi, Linda Loonen, Jos Boekhorst, Houcheng Li, Lingzhao Fang, Peter W. Harrison, Wassim Lakhal, Jerome Lluch, Alexey Sokolov, Sarah Djebali, Andrea Rau, Elisabetta Giuffra, Jerry Wells

## Abstract

This study investigates the impact of maternal gestation diets with varying fiber contents on gene expression and chromatin accessibility in fetuses and piglets fed a low fiber diet post weaning. High-fiber maternal diets, enriched with sugar beet pulp or pea internal fiber, were compared to a low-fiber maternal diet to evaluate their effects on liver and muscle tissues. The findings demonstrate that maternal high-fiber diets significantly alter the chromatin accessibility, predicted transcription factor activity and transcriptional landscape in both fetuses and piglets. A gene set enrichment analysis revealed over-expression of gene ontology terms related to metabolic processes and under-expression of those linked to immune responses in piglets from sows given the high-fiber diets during gestation. This suggests better metabolic health and immune tolerance of the fetus and offspring, in line with the documented epigenetic effects of short chain fatty acids on immune and metabolic pathways. A deconvolution analysis of the bulk RNA-seq data was performed using cell-type specific markers from a single cell transcriptome atlas of adult pigs. These results confirmed that the transcriptomic and chromatin accessibility data do not reflect different cell type compositions between maternal diet groups but rather phenotypic changes triggered by the critical role of maternal nutrition in shaping the epigenetic and transcriptional environment of fetus and offspring. Our findings have implications for improving animal health and productivity as well as broader implications for human health, suggesting that optimizing maternal diet with high-fiber content could enhance metabolic health and immune function in the formative years after birth and potentially to adulthood.

## Introduction

Several studies in humans and animals have reported the benefits of optimizing maternal diets for the health of mothers and offspring via a modulation of the composition of maternal gut microbiota (Lu et al. 2024). Dietary fibers can be considered key ancestral compounds that regulate macronutrient availability and shape the gut microbiota ecology, by serving as substrates for fermentation (Makki et al. 2018). Maternal diet formulations enriched in fiber content have been investigated in farm species in the context of sustainable animal production. Recent studies in pigs have shown that both the amount and sources of dietary fiber in maternal diets influence gut health and modulate immune responses, contributing to performance and health benefits to the sow and offspring (B. Liu et al. 2021; Huang et al. 2023; Sun et al. 2023).

The epigenetic modification of the host by commensal microbes in the gut represents a broad and fundamental level of regulation during early life development, homeostasis, and disease (Woo and Alenghat 2022). While many of the different donor substrates required by epigenetic-modifying enzymes can be generated from host-intrinsic pathways, several of the biological compounds synthesized by microbiota activity serve as epigenetic substrates, cofactors or regulators of epigenetic enzyme activity. Microbiota-sensitive epigenetic changes include modifications to the DNA or histones, as well as regulation of non-coding RNAs, and have been described in both local intestinal cells and in peripheral body tissues (Woo and Alenghat 2022).

Among microbiota metabolites, short-chain fatty acids (SCFA) are epigenetically-relevant molecules that likely play major roles in mediating the beneficial effects of dietary fiber on the host gut and immune system (Woo and Alenghat 2022; van der Hee and Wells 2021). SCFA are exclusively produced by the colon microbiota through fermentation of complex non-digestible carbohydrates and fiber (van der Hee and Wells 2021). In humans and pigs the most abundant SCFA in the mammalian intestinal lumen are acetate, propionate, and butyrate, which are typically found in pig feces at molar ratios of around 60-70% for acetate, 15 to 25% for propionate and 10-15% for butyrate, but can vary depending on the diet, age and health status and microbiota composition. The source of dietary fiber has a large effect on SCFA production and absorption in the hindgut of pigs due to their microbial fermentation in the caecum and colon. A recent study in pigs with several different sources of dietary fiber showed sugar beet pulp (SBP) to produce the highest amount of total SCFA in the feces (Bai et al. 2021). Butyrate is preferentially metabolized by colonocytes via oxidative phosphorylation and serves as an important energy source. The SCFA that are not metabolized by colonocytes enter the blood capillaries, reach the liver via the portal vein, and those not cleared by the liver enter systemic circulation. In germ free (GF) mice, the addition of SCFA partly restored histone H3 and H4 acetylation and methylation patterns in colon, liver, and adipose tissue, which was observed by the colonization of GF mice with fecal microbiota from a conventional mouse, underling the potential of these molecules to drive epigenetic effects at different body sites (Krautkramer et al. 2016).

Research directly addressing the ability of microbial metabolites to have epigenetic effects in the developing fetus are limited. Most studies focus on the roles of SCFA within the maternal gut and their systemic effects rather than their direct transfer to the fetus. In a human cohort study, a correlation was found between the acetate levels in cord blood and maternal blood suggesting that maternal SCFA are likely to cross the placenta and therefore influence fetal SCFA levels (Hu et al. 2019). Thorburn et al. demonstrated that high-fiber or acetate-feeding of pregnant mice mitigated allergic airways disease in the offspring, independent of the offspring microbiota (Thorburn et al. 2015). These findings, among others, suggest that the fetus is exposed to maternal metabolites, some of which can influence the immune system.

In this study we assessed how maternal diets with different fiber contents affect the chromatin accessibility and gene expression landscape of pig fetuses and post weaned piglets. Our hypothesis was that SCFA from the sow would transfer to the fetal bloodstream via placental transport and trigger epigenetic modifications detectable in peripheral organs, and that these epigenetic effects could still be detected in post-weaned piglets. The tested high fiber diets contained sugar beet pulp (SBP) or pea internal fiber, which have been previously shown to stimulate the formation of SCFAs in the pig intestine (Jha and Leterme 2012), as well as a low fiber control diet. The targeted tissues were liver and muscle, whose physiology is profoundly affected by the intestinal microbiota (Pabst et al. 2023; Chew et al. 2022). Both tissues are known to be metabolically modulated by SCFA and host resident immune cells, especially the liver (Kulle et al. 2022).

## Results

### Experimental design and sample collection

Groups of seven sows were fed diets differing in the source of non-digestible fiber content during two successive gestation periods (LF: low fiber diet; HFB: high fiber diet containing SBP; and HFP: a high fiber diet containing pea internal fiber). A schematic representation of the experimental design, timeline, and sample collection points is shown in Fig 1. For each sow, multiple tissue samples from two males and two females were collected from piglets born after the first pregnancy or fetuses 70 days post fertilization (70 dpf) in the second pregnancy.

**Figure 1.**
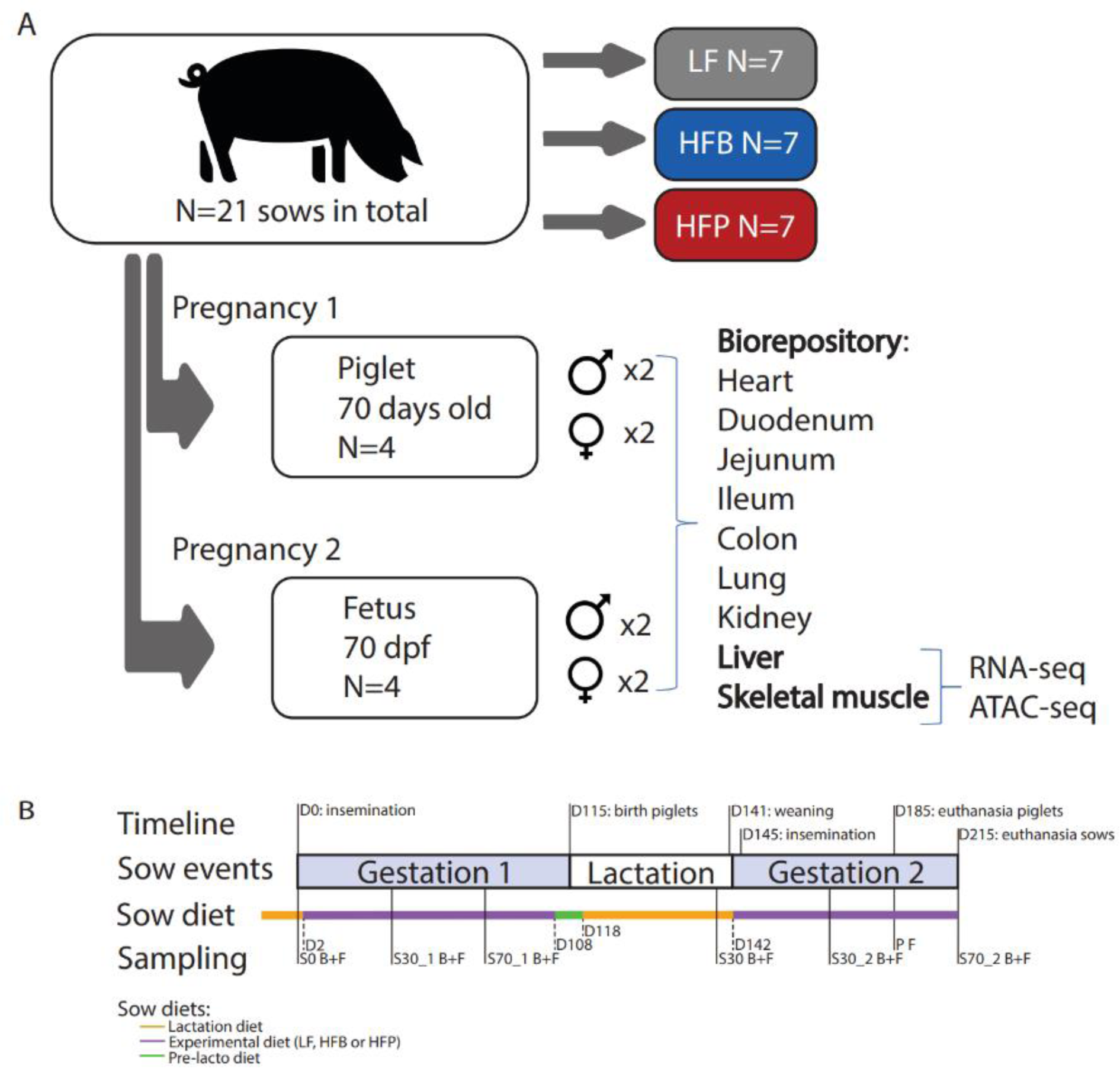
Schematic representation of the experimental design. A) Groups of seven sows were fed one of three diets during two successive pregnancies: low fiber (LF); high fiber diet containing sugar beet pulp (HFB); or high fiber containing pea internal fiber (HFP). Multiple tissue samples were collected from 2 male and 2 female piglets (first pregnancy) and fetuses (second pregnancy) on day 70 post fertilization (dpf). Gene expression (RNA-seq) and chromatin accessibility (ATAC-seq) data were subsequently generated for liver and skeletal muscle tissues. B) Experimental timeline, including times of sampling, sow diet changes, and sow events such as gestation and lactation. For composition and chemical composition of experimental diets and days of diet changes, see Suppl. Tables 1 and 2. S indicates samples taken from sows, P from piglets, and the day when samples were taken is indicated in the “Sampling” row (e.g., S30 indicates day 30 of a sow’s lactation period). B+F means blood + fecal sample, and F feces.

### Differences in maternal diets do not significantly alter the weight of fetuses and piglets

We first sought to establish whether differences in the fiber composition of maternal diets had an impact on the weight of fetuses and piglets. We found no significant effect of maternal diet on fetal weights after accounting for sex, parental breeds, the position *in utero*, and a random litter effect (*n*=80; ANOVA Type III test, *P* = 0.69; Suppl. Figure 1, Suppl. Table 3).

We performed a similar analysis for all piglets born (*n*=392) as well as only those piglets that survived post-weaning and were fed by their birth mother (*n*=253). In line with previous findings in a similar experiment (B. Liu et al. 2021), an ANOVA Type III test accounting for maternal breed, parity, litter size, batch, and a random litter effect showed no significant effect of maternal diet on birth weight (*P*=0.13 for all piglets born; *P*=0.36 for post-weaned piglets). With respect to weaning weight, an ANOVA Type III test accounting for maternal breed, parity, litter size, round, weaning age, number of weaned piglets, and a random litter effect further showed no significant effect of maternal diet (*P=*0.54). Similar results were found for piglet weight gain between weaning and birth, or between weight weaning and sacrifice (Suppl. Figure 2, Suppl. Table 4).

### Impact of different maternal diets on the microbiota and SCFA levels in sows and piglets

Maternal diets induced a significant difference in the composition of the sow fecal microbiome at all sampled time points after day 0 (Figure 2, day 30 of 1st gestation, RDA, *P* ≤ 0.002), but did not lead to a significant change in SCFA measured in both feces and serum (Suppl. Figure 3). However, all fecal SCFA were higher in the HFB group compared to the LF diet, except at day 30 during lactation when all sows were placed on the same lactation diet. A similar trend was observed in the HFP group for acetate and propionate, with the exception of day 30 during the 1st lactation. Fecal butyrate did not show the same trend for HFP, and averages were higher compared to the LF group only during the second gestation. In serum no trends in serum concentration were apparent. Several factors could contribute to this discrepancy, including altered liver metabolism resulting in clearance of SCFA entering the circulation and the timing of sample collection relative to SCFA production and uptake in the colon. Moreover, concentrations of SCFA in the portal vein are known to be substantially higher than in the arteries and likely even higher in the veins leaving the colon to join the portal vein, as indicated by our SCFA measurements in the iliac veins (Suppl. Figure 4).

**Figure 2.**
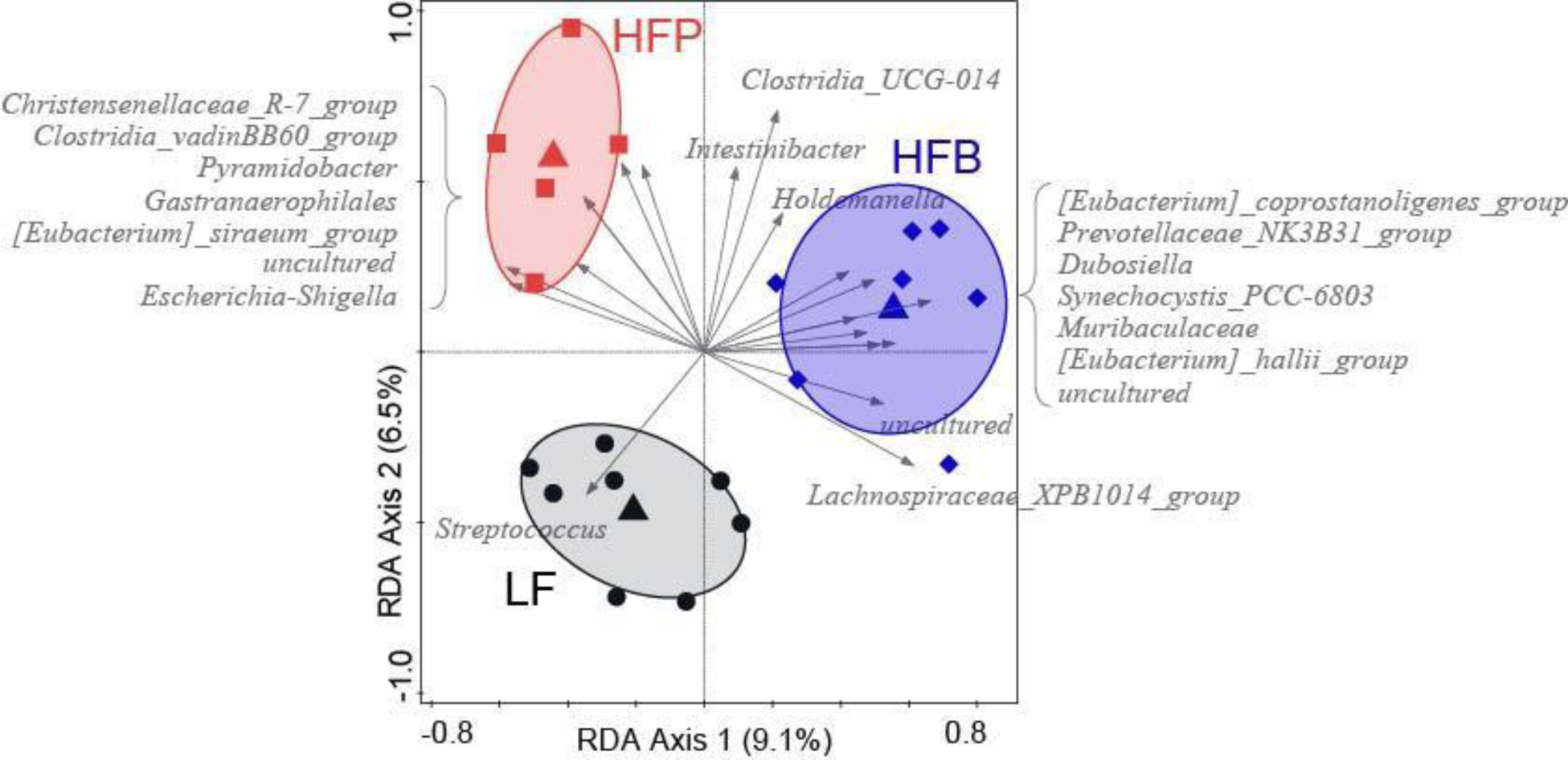
First two axes of the RDA of sow fecal microbiota composition on day 30 of the first gestation, according to the sow dietary group. Relative genus-level abundances were used as response variables and diets as the explanatory variable. Triangles indicate centroids for each diet group, and circles represent individual sows. Gray arrows are the 20 best-fitting genera. Ellipses show the 66% quantile of the approximated 2D-normal density distribution function for each diet group. Colored labels indicate diets: HFP (red), high fiber pea; HFB (blue), high fiber beet pulp; LF: (grey), low fiber.

In piglets at 10 weeks of age, the microbiota composition was not significantly linked to the maternal diet (RDA with genus-level microbiota composition, *P* > 0.1, explained variation 4.35%). Only the link between genus-level microbiota and butyric acid concentration was significant after accounting for the sow effect (Suppl. Figure 5). We also observed significant differences in SCFA levels, notably acetic and propionic acid, in piglet feces at 10 weeks of age between the two high fiber maternal diets (Suppl. Figure 6). However, for both of these SCFA, only the HFP diet group showed a significant increase, while the HFB diet group displayed similar levels as the LF diet group.

### Omics data analyses across tissues and developmental stages

Transcriptome and genome-wide chromatin ATAC-seq profiling were performed to gain insights into the effects of different maternal diets on the functional genome of liver and skeletal muscle of fetuses and piglets.

#### Transcriptome and genome-wide chromatin profiling data

We note that all samples corresponding to piglets from a sow whose diet was erroneously switched during gestation and corresponding to fetuses from a sow who received an antibiotic treatment during the experiment (see Materials and Methods) were removed from the analyses described below.

A total of *n*=304 RNA-seq libraries were generated from the muscle and liver tissues of 160 fetal and 144 piglet samples respectively. RNA-seq libraries were directionally sequenced at a coverage depth of ∼2 × 70M paired-end reads per biological replicate. After removal of an additional 8 samples identified after quality control as outliers with a much lower average number of reads than other samples, RNA-seq analyses were further conducted on *n*=296 samples (Suppl. Table 5). Transcriptome quantification identified on average 17,067 expressed genes (16,776 in fetus liver; 16,817 in piglet liver; 18,986 in fetus muscle; and 15,691 in piglet muscle). Among expressed genes, 72.25% were found to be protein-coding and 21.63% to be lncRNAs (Suppl. Figure 7). These classifications are in line with previous results observed in similar data (Foissac et al. 2019).

A total of *n*=304 ATAC-seq libraries were respectively generated from the muscle and liver tissues of 160 fetal and 144 piglet samples and were sequenced twice at a depth of coverage of ∼2 × 50M reads per biological replicate. Quality controls performed on raw and aligned reads identified an additional 29 samples as outliers that failed quality control checks and were removed from further analyses. Subsequent ATAC-seq analyses were performed on a set of *n*=275 samples (Suppl. Table 5). Genome-wide chromatin profiling revealed 245,471 consensus peaks. Read distribution profiles around TSS (transcription start sites) confirmed an enrichment upstream and downstream TSS, as expected (Suppl. Figure 8).

Principal component analysis (PCA) of normalized ATAC-seq and RNA-seq data (Suppl. Figure 9) revealed a strong separation of samples in both assays between each combination of tissue (skeletal muscle, liver) and developmental stage (fetus, piglet). Both chromatin accessibility and transcriptome data were first structured by tissue (axis 1; 31.06% and 23.3% explained variability in ATAC-seq and RNA-seq, respectively) and then by developmental stage (axis 2; 11.55% and 13.91% explained variability in ATAC-seq and RNA-seq). No significant sample separations were identified when considering combinations of tissue (liver, muscle), developmental stage (fetus, piglet) and sex (Suppl. Figures 10 and 11).

#### Integrative analyses reveal inter-relationships between chromatin accessibility and gene expression

We capitalized on the matched multi-omic data generated within our experimental design to perform an integrative analysis of the chromatin accessibility and transcriptome data to better understand the underlying common relationships and assess the agreement between each omics layer. Rather than estimating individual correlations among expression and TSS accessibility at a gene level, we instead focused on a multivariate assessment of shared signal. We focused on a pairwise assessment of the relationships between matched omics datasets using a coinertia analysis. We found significant positive associations between RNA-seq and ATAC-seq data for piglet muscle (RV=0.46, *P*=0.02) and liver (RV=0.54, *P*=0.01) as well as for fetus liver (RV=0.51, *P*=0.03), although no significant association was found for fetus muscle (RV=0.36, *P*=0.27). This suggests a global correspondence between the RNA-seq and ATAC-seq data that provides a comprehensive regulatory landscape of protein-coding genes and chromatin accessibility, particularly in piglets.

### Functional gene sets, but not individual genes, show altered expression between maternal dietary groups

We detected no differential expression at the individual gene-level between any pair of maternal diets, regardless of tissue or developmental stage (Suppl. Table 6). Thus, we performed a gene set enrichment analysis (GSEA) on the RNA-seq data to identify significantly altered biological processes and pathways between pairs of the dietary groups.

The results of the HFB vs. LF comparison (Suppl. Tables 8-11) are visualized in Figure 3 for the piglet stage. Firstly, immune-related processes in the gene ontology (GO) term clusters for immune response and immune cell activation and differentiation were generally under-expressed in both liver and muscle (Figure 3A). Secondly, in the liver there was a higher expression of genes related to metabolic processes, carboxylic acid transport, and mitochondrial activity (Figure 3B). Thirdly, in the muscle there was a higher expression of genes in GO terms involved in muscle structure and function, transcriptional regulation, RNA processing and protein catabolic processes (Figure 3B). In the 70 dpf fetuses, the two tissues differed by a predominant overexpression in liver and a predominant underexpression in muscle (Suppl. Tables 8 and 9). Both tissues shared overexpression of genes in GO term clusters related to RNA processing, DNA replication, cell division and extracellular matrix, consistent with increased organ development and morphogenesis, whereas underexpression occurred for genes related to metabolic reprogramming and, particularly in muscle, mitochondrial biogenesis. Interestingly, genes in GO terms related to transcriptional regulation and epigenetic modifications (e.g. chromatin organization and histone modifications) were found to be underexpressed predominantly in muscle (Suppl. Table 9). The HFP vs. LF comparison yielded similar results at both stages. The expression of genes linked to protein synthesis (RNP complex, translation and cytosolic ribosome) was slightly higher in piglet and fetus muscle compared to the HFB vs. LF comparison (not shown).

**Figure 3.**
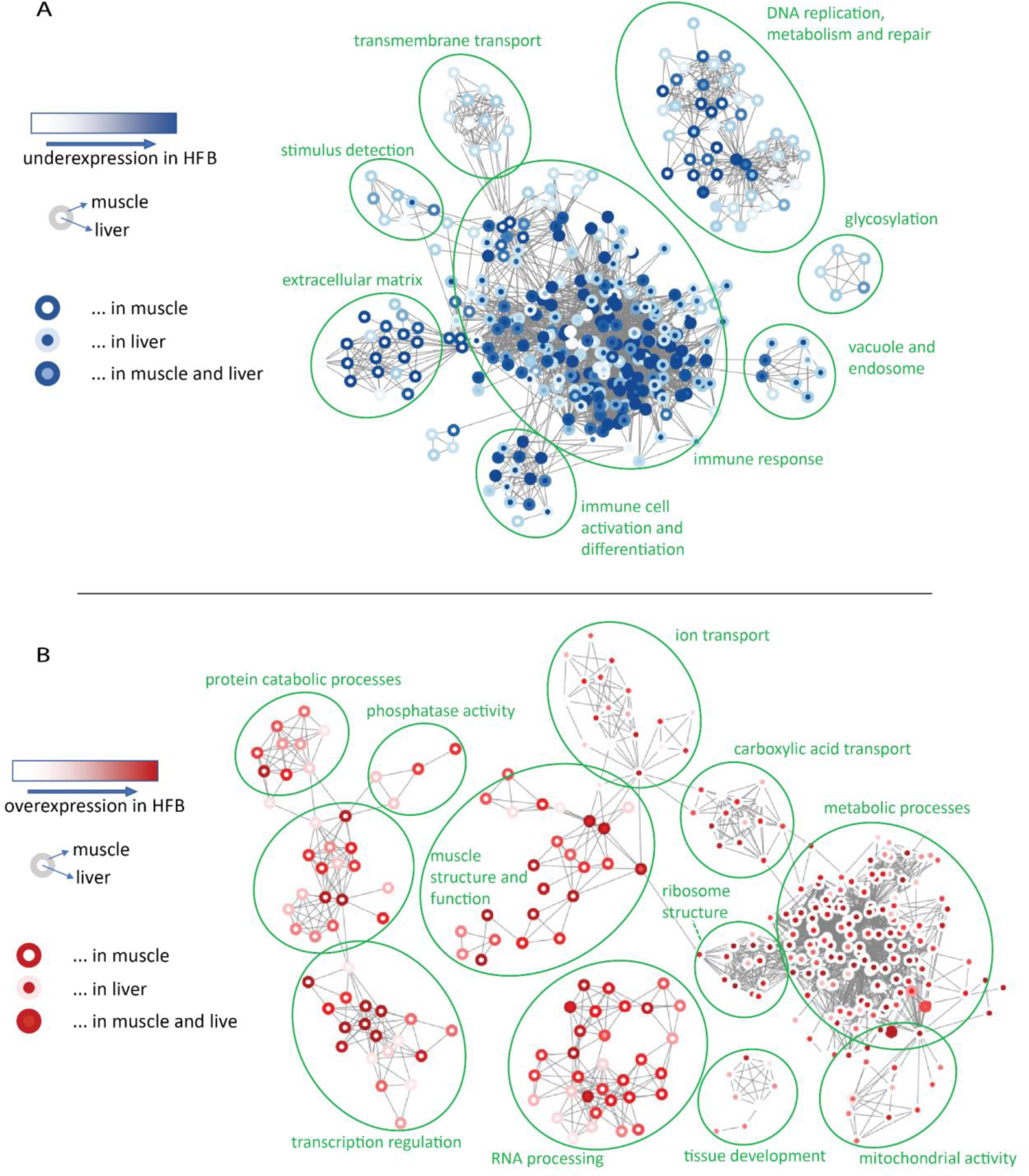
Visualization of RNA-seq GSEA results in piglet liver and muscle for gene ontology (GO) terms that are underexpressed (A, shades of blue) or overexpressed (B, shades of red) in the HFB diet relative to LF. Nodes are GO terms, edges connect GO terms with 50% or more overlap in assigned genes. Node border and fill color reflect significance as determined with ErmineJ (see details in Methods). Green labels indicate the common denominator of the encircled GO terms, as determined through manual inspection. Visualization was performed in Cytoscape.

### Different maternal diets have no influence on the cell composition of liver and muscle

More insights into the GSEA results were obtained by investigating the changes in cell type compositions between the HFB, HFP and LF groups. To this end, a deconvolution analysis of the bulk RNA-seq data was performed in liver and muscle at both developmental stages using cell-type specific markers from a single cell transcriptome atlas of adult pigs (Chen et al. 2023).

As a first step, we explored how the relative proportions of cell types varied across the two developmental stages (Suppl. Figure 12AB). As expected, in liver samples hepatocytes emerged as the dominant cell type at both stages, closely followed by macrophages. Both cell types exhibited notable variability across samples, as a reflection of individual differences. In muscle samples, type IIx myonuclei, type I myonuclei, and fibro-adipogenic progenitors were consistently abundant across both stages. The missed representation of functional type IIa.b myonuclei in 70 dpf fetuses is consistent with previous findings on muscle ontogenesis for pigs (Picard et al. 2002). To further pinpoint the developmental effects on cell components, we extracted three cell types with different cell component levels to perform Student’s t-test (Suppl. Figure 12CD). In the liver, hepatocytes and macrophages had higher enrichment in fetal samples; in contrast, mesenchymal stem cells were notably more abundant in the piglets. In muscle, fibro adipogenic progenitors and smooth muscle cells were significantly enriched in fetal samples, and Type IIa.b myonuclei had an opposite enrichment. These findings indicated that, as expected, cell components of both common cell types and rare cell types undergo significant alterations in response to individual developmental stages.

Subsequently, we pinpointed the effects of maternal diet on these two tissues. Cell types in each tissue were stably distributed across different diets (Figure 4). We also performed Student’s t-test within several cell types in the three diets and found no significant differences between them regardless of the tissue, indicating that changes in maternal diet have no influence on cell components (Suppl. Figure 13).

**Figure 4.**
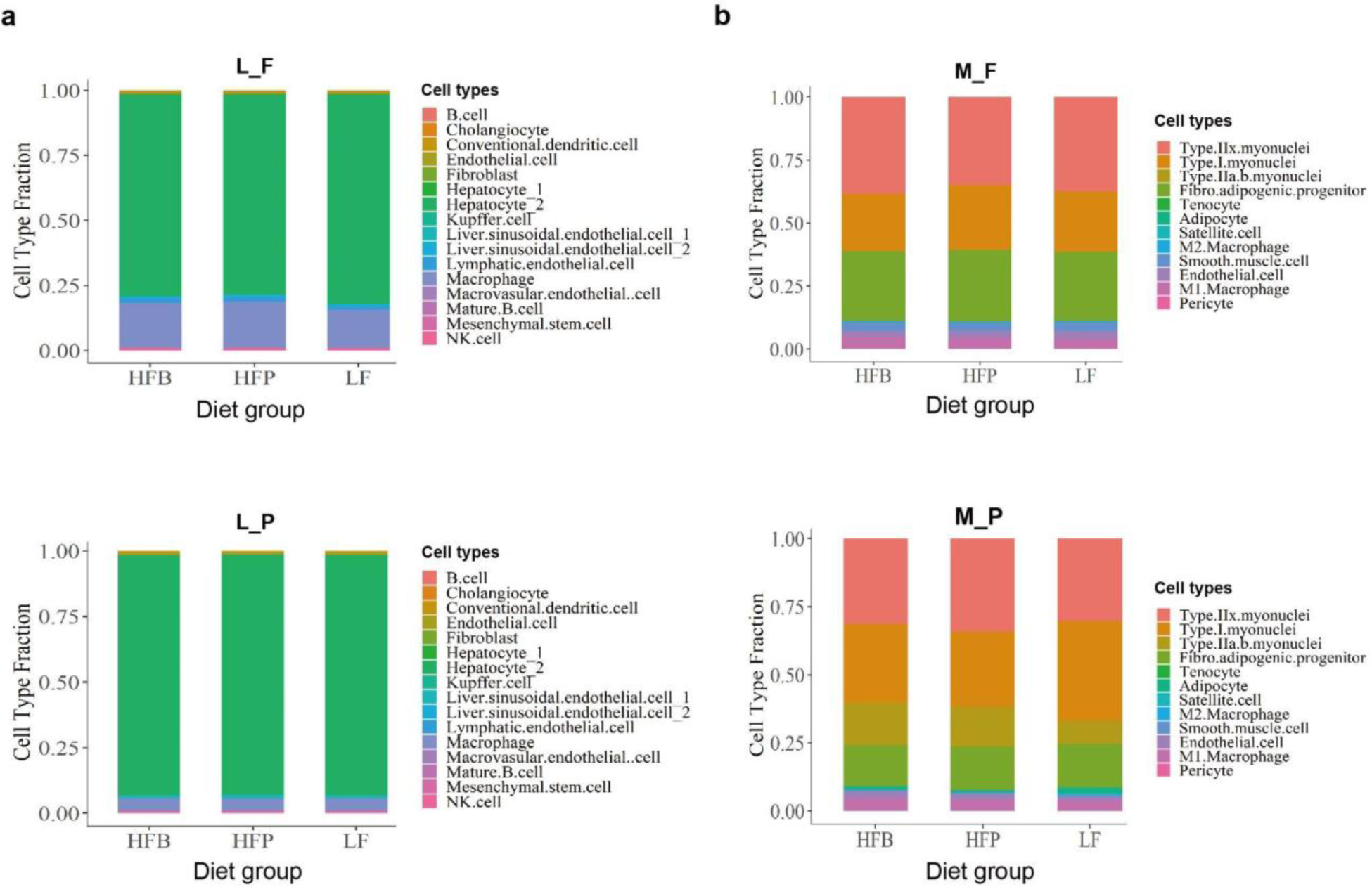
Deconvolution results according to different maternal diets. (A) Average cell component distribution of fetal (L_F) and piglet (L_P) liver samples in three diets. (B) Average cell component distribution of fetal (M_F) and piglet (M_P) muscle samples in three diets.

### Differences in maternal diets significantly affected chromatin accessibility, mainly in piglets

For ATAC-seq data, we first performed differential analyses to identify diet-specific differentially accessible regions for each combination of tissue and developmental stage. As for RNA-seq (Suppl. Table 7), this analysis accounted for maternal and paternal breed, sex, and a random litter effect.

While chromatin accessibility differed considerably between sexes, sow and sire breeds at both the fetus and piglet stages (Suppl. Table 7), we detected a low number (706; FDR < 0.05) of diet-specific differentially accessible chromatin peaks corresponding to a total of 659 differential peaks between HFB and LF, 32 between the HFB and HFP diets, and 15 between HFP and LF. Most of these peaks were identified in piglet muscle for the HFB vs LF comparison, and were mainly over-accessible in the HFB diet group (Suppl. Table 7). A classification of the consensus peaks into annotation categories indicated that 55.5% of differential peaks were localized in gene intronic regions. The diet-specific differentially accessible regions were found to be preferentially located in introns and the promoter-transcription start site (TSS +/− 1kB) of genes (Suppl. Figure 14). To assess any potential sex-specific diet effects, we repeated the analysis with a diet x sex interaction term. Although detection power for such interactions is likely low given the experimental design, we note that no significant effects or trends towards significance were observed.

### Maternal diets triggered differential activity of transcription factors in both fetuses and piglets

More resolutive insights into the pattern of ATAC-seq profiles were obtained by transcription factor (TF) footprint analyses, i.e. by considering all 245,471 consensus peaks to identify active TF binding sites and by comparing TF activity across each combination of tissue and developmental stage for all pairwise maternal diet comparisons.

This analysis identified 21 and 30 transcription factors (TFs) with differential activity in fetuses and piglets, respectively (Figure 5). TF activities were mostly upregulated in fetal liver, while the opposite occurred in fetal muscle. At the piglet stage, only a few TFs displayed a significant (downregulated) pattern in the liver, while muscle was characterized by an intense downregulation of several other TFs. Most effects were found in the pairwise comparisons of HFB vs. the low fiber control (LF), while HFP vs LF showed few effects; however, significant effects were also found in the comparison between HFB and HFP. In total, five TFs were common between stages, namely the three subunits of the NF-Y nuclear factors, DUX (Double Homeobox) and ATF1 (activating transcription factor 1), with the most significant Z-scores found for NFY and DUX. In fetuses (Figure 5A), both the HFB vs. LF and HFB vs. HFP comparisons showed increased activity in liver and decreased activity in muscle of NF-Y. At the piglet stage, and only for the HFB vs. LF comparison, the three NF-Y subunits switched to downregulation at the piglet stage in liver while in muscle only NFY-A and NFY-C were lowly significantly upregulated (Figure 5B). For DUX, the downregulation in fetal muscle was more intense for the HFB vs. LF comparison and upregulation was found in fetal liver, while no significance was observed for both tissues at the adult stage.

**Figure 5.**
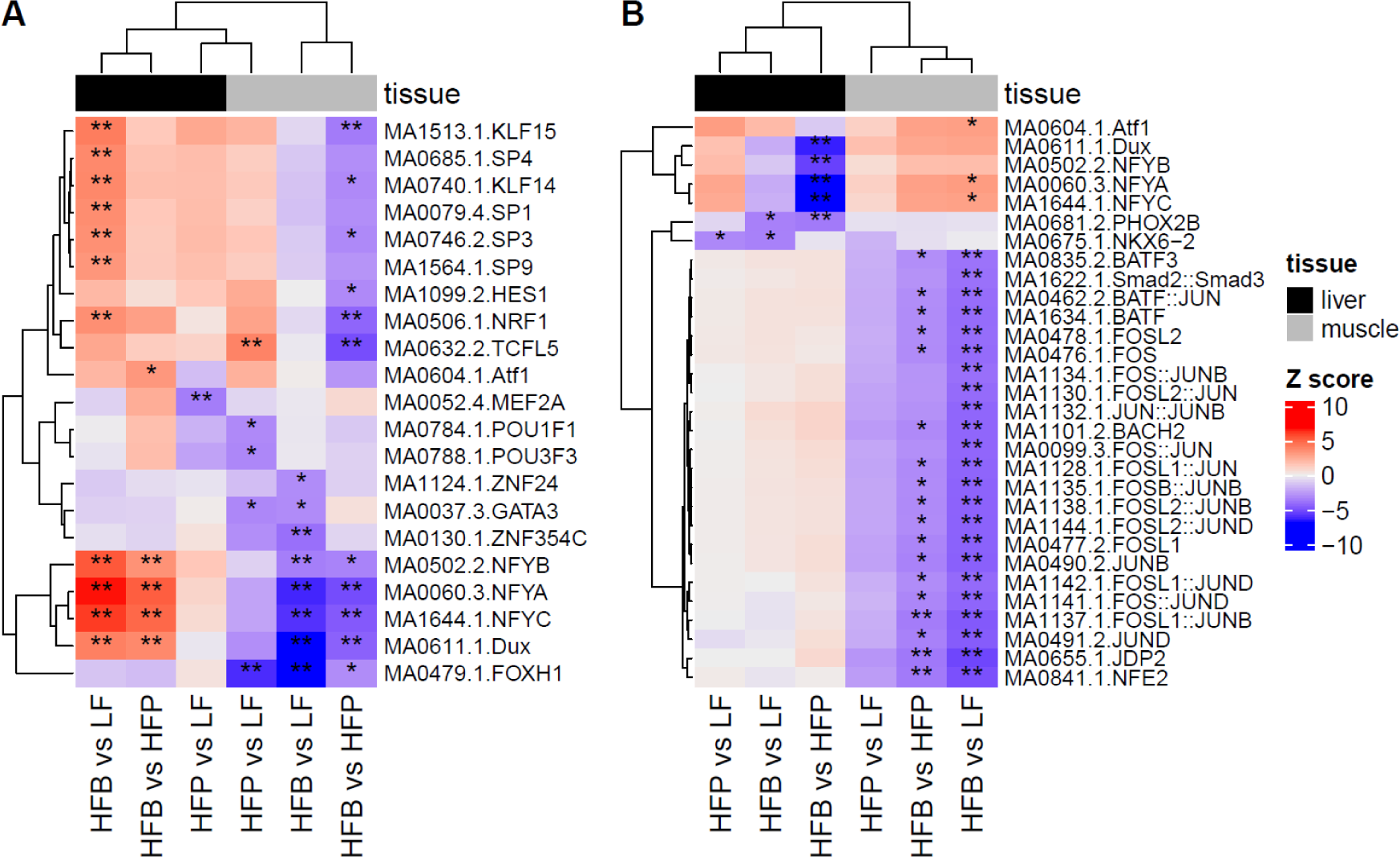
Heatmap of differential activity scores of transcription factor (TF) motifs in (A) fetuses, for 21 TFs and (B) piglets, for 30 TFs. Differential activity scores correspond to Z-scores. Dendrograms on rows and columns correspond to a hierarchical clustering with Euclidean distance and complete linkage. Stars represent the significance level for differential TF activity (“*“: FDR < 10%, “**”: FDR < 5%).

## Discussion

Given the increasing interest in the role of microbial metabolites and especially SCFA in regulating gut barrier defense and immunity in humans and model animals, we sought to study the maternal effects of fermentable fibers on the functional genome of the developing fetus and offspring, targeting the liver and muscle tissues (Figure 1).

As expected, the microbiota composition of sows in the HFP and HFB groups showed distinct clustering away from the LF group, with significant differences based on the fiber content and type in the maternal diet at all sampled time points (Figure 2). These findings align with previous research indicating that dietary fibers modulate gut microbiota composition, thereby impacting health outcomes (Makki et al. 2018; B. Liu et al. 2021). Furthermore, the distinct clustering of microbiota compositions between different high-fiber diets suggests that the type of fiber, in addition to the amount, plays a crucial role in shaping gut microbial ecology.

Intestinibacter were more closely associated with the HFP group, suggesting it may play a role in fermenting pea fiber, whereas Lachnospiraceae_XPB1014_group were more closely associated with the HFB group, indicating its potential role in fermenting beet pulp fiber. Both genera are known to be involved in the colonic fermentation of complex carbohydrates and the production of SCFA (Gao et al. 2022; Tsukuda et al. 2021). The presence of these taxa is consistent with the increased levels of SCFA levels, particularly acetic and propionic acids, that we found in sow feces (Suppl. Figure 3) and with previous findings in pigs fed with diets containing various fiber sources, such as wheat bran, corn bran, and sugar beet pulp (Bai et al. 2021). In our data, SCFA levels were not mirrored in the sow serum (Suppl. Figure 3). However, a recent systematic review of the effects of dietary fiber on SCFA in human feces has shown that dietary fibers do not produce a significant increase in SCFA levels. SCFA production was concluded to be influenced by type and dose of fiber and potentially the microbiota composition (Vinelli et al. 2022).

Subsequent omic analyses in the liver and muscle of 70 dpf fetuses and piglets provided robust evidence for epigenetic effects induced by high fiber contents in the maternal diet. Overall, our results are consistent with the documented epigenetic effects of SCFA on immune and metabolic pathways in adults and with previous evidence that maternal microbiota-derived SCFA can be transferred across the placenta to embryos (Kimura et al. 2020). Importantly, the deconvolution analysis based on bulk RNAseq data allowed us to verify that these analyses are not biased by potential misleading effects due to different cell type compositions between diet groups (Figure 4).

GSEA results are particularly striking in piglets, as the sows alone received the high-fiber or control diets prior to parturition and lactation. Piglets from sows fed with the high fiber diets had reduced expression of genes associated with immune response and immune activation pathways in both liver and muscle (Figure 3A), suggesting reduced inflammation, improved metabolic health and enhanced immune tolerance. This is consistent with several studies demonstrating the anti-inflammatory effects of SCFA in rodents and humans (Visekruna and Luu 2021; Liu et al. 2023). For example, propionate and butyrate are known to suppress immune responses in macrophages and monocytes via inhibition of HDAC activity (X.-F. Liu et al. 2023). The overexpressed GO term clusters in liver (metabolic processes, carboxylic acid transport and mitochondrial activity) (Figure 3B) are consistent with the central role of liver in glucose metabolism and lipid metabolism, which are of particular interest in humans for interventions in liver and metabolic diseases (Canfora, Jocken, and Blaak 2015; Morrison and Preston 2016). In muscle, the overexpression of genes linked to muscle structure and function suggests improved muscle function and development, in line with the proposed role of SCFA in skeletal muscle energy metabolism and muscle fiber conversion (Yin et al. 2022). The overexpression of genes linked to transcriptional regulation and RNA processing could reflect an overall increase in transcriptional activity, potentially improving protein synthesis and muscle growth, and catabolic processes may be crucial for muscle adaptation and remodeling, especially under varying nutritional conditions.

The TF footprint analysis revealed sharp differences between tissues and stages. In fetuses, TFs showed increased activity in liver and decreased activity in muscle, whereas in piglets most TFs showed little significance in liver and significantly decreased activity in muscle (Figure 5). This last finding is corroborated by the larger proportion of differentially accessible peaks found in piglet muscle (Suppl. Table 7), although none of them overlapped with TF loci and no directly associated differential gene expression was found. Interestingly, the effects at both stages and tissues showed quite clear differences between the two dietary fiber types, consistent with the independent clustering of microbiota species in the HFB and HFP sow groups (Figure 2).

The kruppel-like factors KLF15 and KLF14 play roles in glucose metabolism, with KLF14 also involved in regulating the immune system (Yuce and Ozkan 2024). Their predicted increased activity in fetus liver in the HFB maternal diet group is consistent with over-expression of genes in GO terms for metabolic processes. Similarly, the increased activity in fetal liver of two members of the specificity protein family (SP1 and SP4) is consistent with overexpression of genes in the GO terms linked to morphogenesis and cytoskeleton (Suppl. Table 8). Among the five TFs with significantly different activities at both stages, NF-Y and DUX had the most significant Z-scores for both the HFB and HFP dietary groups in fetal liver and muscle. NF-Y is a highly conserved and ubiquitously expressed heterotrimeric TF composed of NF-YA, NF-YB and NF-YC subunits that binds to CCAAT boxes, common regulatory DNA elements within promoters and enhancers (Oldfield et al. 2019). This complex has been implicated in several physiological functions in different tissues. For example, NF-Y is a key regulator of gluconeogenesis in the liver (Yanjie Zhang et al. 2018) and is involved in stem cell fate decision and regeneration in skeletal muscle (Rigillo et al. 2021). The DUX genes encode a family of transcription factors (DUXA, DUXB and DUXC subfamilies) that are expressed in a short burst during early stages of development and, with the exception of testis and thymus, are silenced in most somatic tissues under normal physiological conditions (De Iaco et al. 2017). Emerging data support a role for DUX4 in the control of viral infection and it has been shown that in muscle cells, DUX4 protein interacts with STAT1 and broadly suppresses the expression of IFNγ-stimulated genes (Spens et al. 2023; Mocciaro et al. 2021).

Overall, these results highlight a pervasive and statistically significant epigenetic effect of maternal diets on the functional genome of fetuses and piglets and have broader implications for studies exploring the role of high fiber content in human health. The observed differences in liver and muscle metabolism as well as immune regulation underscore the importance of early nutritional environments in shaping the epigenetic landscape, with potential long-term implications for offspring health and development. Further research is warranted to elucidate the mechanistic long-term effects of these early dietary influences and their potential use in the development of new nutritional strategies in pig production.

## Materials and Methods

### Animal experiment and sampling

The animal experiment was conducted at Schothorst Feed Research in Lelystad, the Netherlands, and approved by the ethical committee from Wageningen University, the Netherlands, under number VA19-10 (AVD246002015280-1). Groups of *n*=7 parent sows (TN60 or TN70 line; Topigs Norsvin) with similar parity (average parity = 5.5 ± 1), were inseminated with terminal sires (TN Duroc or TN Tempo line; Topigs Norsvin) and then fed one of three diets differing in non-digestible fiber (NDF) content (high fiber beet, HFB; high fiber pea, HFP; low fiber, LF) between day 2 after insemination and day 108 of two successive pregnancies (Figure 1). The detailed composition and chemical composition of each diet is described in Suppl. Tables 1 and 2. All sows were housed together and were identified via ear tag recognized by the trough to receive their assigned diet. Sows received *ad libitum* fresh drinking water and were fed according to a feeding scheme for gestating sows (scheme from Schothorst Feed Research). Two sows died of heat stress during their first pregnancy but were replaced by 2 new sows for the second pregnancy. Following birth, all sows were provided with a commercial lactation diet from day 108 of gestation onwards (Figure 1B). As the diet for one sow (ID n° 115) was erroneously switched during the first pregnancy, her fecal and blood samples and all samples from her piglets were omitted from subsequent analyses. Blood (serum, from jugular vein) and fecal samples (after rectal stimulation) were collected from each sow throughout the experiment (Figure 1B) for measurements of SCFAs and microbiota composition.

Piglets born from the first pregnancy were weaned around day 28 after birth. From each litter, 2 male and 2 female piglets were selected based on average birth and weaning weights. These piglets were housed per pen and received *ad libitum* feed and drinking water. Tissue and fecal samples were collected from *n*=72 weaned piglets at around 10 weeks of age to determine whether any effects of the maternal diet on the epigenome and transcriptome would persist in the offspring to post-weaning age. In addition, birth, weaning, and sacrifice weights were recorded for each piglet.

During the second pregnancy, we collected tissue samples from fetuses 70 days post-insemination (dpi) to compare the effects of the different maternal diets on fetal tissues *in utero*. As one sow (ID n° 173) received an antibiotic treatment during gestation for ethical reason, her fetuses were omitted from all subsequent analyses, leading to a total of *n*=80 fetuses used in the study. After sedation with Zoletil, sows were euthanized using an intracardiac injection of T61®, after which 2 male and 2 female fetuses were taken from the uterus. The most important selection criterion was choosing either male or female fetuses in proximity to the bifurcation of one of the two uterine horns.

In total, small intestine, colon, heart, distal lung, kidney (cortex), liver and skeletal muscle samples were collected for each piglet (*n*=72) and fetus (*n*=80) and snap frozen in liquid nitrogen after dissection. A biorepository of all tissue samples collected in the animal experiment was stored at −80°C at WUR facilities (Wageningen, the Netherlands). Data for individual specimens and all associated metadata (e.g., insemination date, pregnancy length, gestational age at collection for fetuses, birth and weaning weights for piglets, sex, etc) were uploaded to the FAANG data portal (https://data.faang.org/projects/GENE-SWitCH).

In this work, we selected the liver and skeletal muscle as target tissues for functional genomics assays. In total, in the current study we performed functional genomics assays in two tissues (liver, skeletal muscle) at two developmental stages (fetus, piglet), corresponding to a total of *n=*304 samples, with a roughly even balance across the 3 maternal diets.

### Sow and piglet fecal microbiota analysis

To isolate bacterial DNA from sow and piglet feces for microbiota profiling, the Qiagen Powersoil PRO DNA isolation kit was used. Isolation was performed according to the manufacturer’s manual. Two cycles of 40 seconds each at 4.0m/s were run using the beat beater (FastPrep). Elution was done with 50µl solution C6. After measuring the concentration by Qubit the DNA was stored at −20°C. When all samples were isolated, DNA was diluted to 50µl of 20ng/µl in MQ, after which the samples were sent to Genewiz (Germany) for sequencing.

The *QIIME* data analysis package (version 1.8) was used for 16S rRNA data analysis (Caporaso et al. 2010). The forward and reverse reads were joined and assigned to samples based on barcode and truncated by cutting off the barcode and primer sequence. Quality filtering on joined sequences was performed and sequences that did not fulfill the following criteria were discarded: sequence length < 200bp, no ambiguous bases, mean quality score ≥ 20. Sequences were then compared to the reference database (RDP Gold database) using the UCHIME algorithm to detect chimeric sequences, and chimeric sequences were then removed. Only effective sequences were used in subsequent analyses. Sequences were grouped into operational taxonomic units (OTUs) using the clustering program *VSEARCH* (version 1.9.6) against the Silva 119 database pre-clustered at 97% sequence identity. The Ribosomal Database Program (RDP) classifier was used to assign a predicted species-level taxonomic category to all OTUs at a confidence threshold of 0.8. Sequences were rarefied prior to the calculation of alpha and beta diversity statistics.

### Sow serum SCFA measurements

SCFAs were derivatized and measured using LC-MS. For derivatisation, 40µl of thawed serum was used. We added to the serum 5µl SCFA reference mix (10mM stock, Sigma, prepared in different working solutions) for calibration curves, or 5µl MQ to the actual samples. Next, a 3µl 4x diluted 13C-labeled SCFA mixture (Sigma) was added. The samples were diluted 1:3 with ice-cold pure acetonitrile (ACN, Sigma), and stored on ice. After 15 minutes, the samples were vortexed and centrifuged for 20 minutes at 4°C and 12500g. One hundred microliters of the supernatant was mixed with 100µl 200mM 3-Nitrophenylhydrazine hydrochloride (3NPH in 50% ACN) and 100µl 120mM N-(3-dimethylaminopropyl)- N’-ethylcarbodiimide hydrochloride (EDC in 50% ACN with 6% pyridine) (both Sigma) in a low-binding Eppendorf tube. These components were carefully mixed and incubated in a thermomixer (Eppendorf) at 40°C for 30 min at 400 rpm. The tubes were put on ice for 1 minute after incubation. One hundred microliter MQ was added, and the samples were centrifuged for 5 minutes at 4°C at 12500g. One hundred microliter of the supernatant was transferred to an UPLC vial containing an insert (BGB), after which they were stored cold until measurement. The samples were measured using a Shimadzu LC-20AD XR UPLC system coupled to LCMS-8050 triple quadrupole MS, using a Phenomenex Kinetex C18 column (50 x 2.1mm, 1.7 µm). Flow rate was 0.6 mL/min and 10 µL of sample was injected onto the column. The method used MQ water with 0.1% formic acid as mobile phase A and Acetonitrile with 0.1% Formic Acid as mobile phase B. The following gradient was used: 10% B at 0 min; 20% B at 4.0 min; 100% B at 4.1-7.5 min and 10% at 7.6-15 min. Water samples were measured in between batches of samples, and calibration curves were measured at the beginning and end of each sample set.

### Sow and piglet fecal SCFA measurements

Approximately 500g of fecal sample was added to MQ at a ratio of 1:4 (w/v), and thoroughly mixed by vortexing at 2100 rpm for 15 min. The mixture was centrifuged for 5 min at 4°C and 14000 rpm, and the supernatants were filtered through a 0.22 μm organic filter. An aliquot (100 μL) of the supernatant was acidified by adding 50 μL of a solution of 0.3 mol/L HCl and 0.9 mol/L oxalic acid containing 0.45 mg/mL 2-ethylbutyric acid as the internal standard. This mixture was then subjected to SCFA measurement by a Shimadzu GC-2014, equipped with a Flame-ionization detector (FID), a capillary fatty acid-free Stabil wax-DA column (1 μm × 0.32 mm × 30 m) (Restek) and a split injector. The injection volume was 0.5 μL and the carrier gas was nitrogen. Temperature of the injector and detector were 100 and 250°C, respectively. The temperature profile started at 100 °C, increased to 172 °C by 10.8 °C/min, then to 200 °C by 50 °C/min, and lasted for 1 min. Standard solutions of SCFAs (acetate, propionate, butyrate, valerate, isobutyrate, and isovalerate) were prepared and used for identification and quantification. The results were processed using Chromeleon 7.2.10 (Thermo Fisher Scientific).

### PolyA+ RNA isolation and sequencing

RNA was isolated from liver and skeletal muscle tissues in fetuses and piglets using the RNA isolation protocol uploaded on the [FAANG Data Portal](https://data.faang.org/api/fire_api/experiments/WU_SOP_GENESWITCH_WP5_RNA_ISOLATION_20211108.pdf). Briefly, tissues were dissociated using a Miltenyi GentleMacs, after which the Qiagen RNeasy kit was used for RNA extraction, including an on-column DNase treatment. This method worked well for liver; in muscle, due to initial low RNA yield this approach had to be optimized by increasing the amount of tissue and including a proteinase K treatment of the homogenized tissue. The optimized method enabled a high yield of good quality RNA from all *n*=304 samples as measured by nanodrop. The purified RNA was then sent to iGenSeq (France) for preparation of RNA libraries and sequencing. Sequencing was performed using an Illumina NovaSeq 6000.

Raw data and associated metadata were uploaded on the EBI server (ERP137576) and are available on the FAANG Data Portal (https://data.faang.org/dataset/PRJEB52831). Based on quality control checks, the raw data were found to be of good overall quality, although 8 of the 304 samples were found to have average number of reads much lower than the others and were thus removed from subsequent analyses (Suppl. Table 12).

### RNA-seq data processing and analysis

Gene expression was quantified using the *TAGADA* pipeline (Kurylo et al. 2023), which enables transcript reconstruction and quantification of transcript expression (https://github.com/FAANG/analysis-TAGADA). Briefly, sequenced reads were first mapped to the pig genome reference version 11.1.108 with *STAR* (Dobin et al. 2013), and expression quantification was performed with *StringTie* (Pertea et al. 2016).

Data were normalized as counts per million (CPM) using the trimmed mean of M-values (TMM) approach (Z. Li et al. 2023; Robinson and Oshlack 2010) from the *edgeR* Bioconductor package (Robinson, McCarthy, and Smyth 2010). Principal component analyses (PCA) on log-normalized expression data were used to visualize transcriptome-wide patterns in samples across tissues and development stages. To filter genes with globally weak expression, only genes with a CPM > 10 in at least 10 samples were retained for the analysis. Differential analyses were performed with log-normalized CPM values using the *limma-voom* approach with empirical sample quality weights (R. Liu et al. 2015). Briefly, for each combination of tissue and developmental stage we used per-gene linear models, including sex, sow breed, boar breed (for fetuses only), and maternal diet as fixed effects and sow as a random effect. Contrasts were estimated to perform tests for comparisons of interest, notably between pairs of maternal diets. *P*-values were corrected for multiple testing (Benjamini and Hochberg 1995), and a significance threshold of FDR < 5% was used.

A more generic overview of biological processes impacted by maternal diet was generated by gene set enrichment analysis (GSEA) using *ermineJ (Lee et al. 2005)* on log-fold change values with the receiver operating characteristic (ROC) method. GSEA ranks genes by pertinent signals (e.g., log-fold changes or associated *P*-values) for a given comparison of interest to subsequently identify biological pathways or processes, such as GO terms, that are enriched in highly ranked genes.

### ATAC-seq library preparation and sequencing

Chromatin accessibility was quantified using the Assay for Transposase-Accessible Chromatin with high-throughput sequencing (ATAC-seq) protocol (Buenrostro et al. 2015). Libraries were prepared by Diagenode, according to an optimized protocol on frozen tissues (Corces et al. 2017). Biological replicates were sequenced on an Illumina NovaSeq 6000 (GeT-PlaGe platform) with 150bp paired-end reads and an average (median) depth of coverage of approximately 2 x 49 (42) million paired-end reads per replicate.

Raw data and associated metadata were submitted to the FAANG DCC under the study accession ID ERP138237 and are available on the FAANG Data Portal (https://data.faang.org/dataset/PRJEB53440).

### ATAC-seq data processing and analysis

ATAC-seq data were processed using version 1.2.1 of the nf-core ATAC-seq pipeline (Ewels et al. 2020), found at https://nf-co.re/atacseq. Samples belonging to a given combination of tissue (liver, muscle), developmental stage (fetus, piglet), and maternal diet (HFB, HFP, LF) were considered to represent a single condition for peak calling. Within the pipeline, reads were mapped with *bwa* (H. Li and Durbin 2009) on the pig genome reference version 11.1.108 and peaks were called from the mapped reads of each condition with *macs2* (Yong Zhang et al. 2008). Peaks obtained in each condition were then merged together into consensus peaks, which were annotated relative to genes (either overlapping TSS or according to the closest position) using *HOMER* (Heinz et al. 2010). Mitochondrial peaks were removed as artifacts. Quality control measures flagged a total of *n*=29 samples as being outliers (Suppl. Table 12): (1) weak score for enrichment at gene TSS (*n*=4), which quantifies the relative enrichment of signal in promoter regions that tend to be consistently enriched in accessible chromatin; (2) weak fraction of reads mapped into called peak regions (FRiP score), which assesses the enrichment of transposase insertions in regions of known chromatin accessibility. ENCODE project standards (https://www.encodeproject.org/atac-seq) recommend the use of 30% as an appropriate threshold for FRiP scores; here we have discarded samples with a FRiP < 10% (*n*=8); (3) per-sequence GC content fails (*n*=3); (4) high percentage of trimmed basepairs (*n*=14); and (5) a too high fraction of duplicated reads (i.e. do not PASS fastqc duplication assessment or with a picard duplication level ≥ 0.3; *n*=3).

Non-linear trended efficiency biases were normalized using loess normalization using the *csaw* Bioconductor package (Lun and Smyth 2016). As for the transcriptomic data, PCA were performed on log-normalized accessibility data to visualize genome-wide patterns across tissues and developmental stages. Peaks with globally weak accessibility across samples were filtered, and only those with CPM > 10 in at least 10 samples were retained. Differential analyses were performed for each combination of tissue and developmental stage with log-normalized CPM values using *limma-voom* with empirical sample quality weights (R. Liu et al. 2015), as described for the RNA-seq data above.

Transcription factor (TF) footprint analyses were performed to detect putative TF binding sites associated with each combination of tissue, developmental stage, and maternal diet using the *HINT* computational tool (version 1.0.0) from the Regulatory Genomics Toolbox (Z. Li et al. 2023). To compare changes in TF activity with differential footprinting, *HINT* generates average ATAC-seq profiles around the putative binding sites of a given TF to compare cleavage profiles across groups of interest (here, maternal diets), thus providing insight into binding changes between the two. Briefly, footprint calling was first performed with the *rgt-hint footprinting* command to identify predictive footprints (Salavati et al. 2022). Predicted footprints were then matched to known motifs in the JASPAR database with the *rgt-motifanalysis* matching command (Castro-Mondragon et al. 2022). For some TFs, the JASPAR database includes different variants of motifs, which were removed here. We retained only TFs for which at least 1000 binding sites were predicted for each combination of tissue, developmental stage, and maternal diet. Finally, TF differential activity between diets was assessed using bias-corrected signal.

### Cellular deconvolution analysis

Marker genes of each cell type were collected from liver and muscle tissues in the pig cell atlas (Chen et al. 2023). For each tissue, differentially expressed genes specific to each cell type were identified using the Findmarkers function in the *Seurat* version 4.0.6 package (Hao et al. 2024). Then the top 50 genes with the most significant overexpression were selected (adjusted *P* < 0.05 and average log2-fold change > 0.5) to build the gene expression signature matrix for the cell-type reference set. Subsequently, the *CIBERSORT* version 1.0.4 tool (Newman et al. 2015) was used for cellular deconvolution analysis on *n*=152 and *n*=144 bulk samples using the TPM matrix of signature genes from each cell type in pig liver and muscle, respectively. The number of permutation tests was set to 1,000, and a threshold of *P* < 0.05 was used to determine statistical significance.

### Statistical analyses

To assess the effect of the maternal diet on fetus weight or piglet birth weight, weight gain during suckling, or weaning weight, we fit linear mixed models including a random litter effect using the *nlme* package (Pinheiro and Bates 2000), with type III analysis of variance (ANOVA) tables using the *car* package (Fox and Weisberg 2018). For fetuses, fixed effects were fitted for sex, maternal and paternal breed, position *in utero*, and maternal diet. For piglet birth weights, fixed effects were fitted for round (e.g. batch), maternal breed, parity, litter size, and maternal diet. Additional fixed effects for age at weaning and number of weaned piglets in the litter were further included for piglet weaning weights and for the weight gain from birth to weaning. Analyses were performed for all piglets born (*n*=392) as well as only those that survived post-weaning and were fed by their birth mother (*n*=253).

Redundancy analyses (RDA) were performed with *Canoco* v5.15 (Braak and Šmilauer 2012) using a constrained analysis. For microbiota analyses, genus-level relative abundances were log-transformed after multiplication by 1000 and the addition of a constant of 1, and were subsequently used as response variables. To evaluate the link between piglet microbiome and sow diets while accounting for a sow effect, we performed hierarchical permutation tests (whole-plots freely interchangeable, no split-plot permutations) to permute litters rather than individual piglets among diets. For the analysis of SCFA levels and piglet microbiota composition, we included sow as a covariate.

Coinertia analysis is a multivariate statistical analysis to assess the covariance between two tables of variables collected on the same set of individuals by identifying components that maximize the covariance between the two (Dray, Chessel, and Thioulouse 2003). We performed coinertia analyses on log-normalized counts of expression (RNA-seq) for genes identified with TSS peaks and log-normalized chromatin accessibility (ATAC-seq) for TSS peaks in muscle and liver for piglets and fetuses using the *ade4* package (Thioulouse et al. 2018). For each combination of tissue and developmental stage combination, we estimated the RV coefficient, a multivariate generalization of the squared correlation coefficient. A permutation test based on 99 random permutations of piglets was used to evaluate the significance of each RV coefficient.

Unless otherwise noted, statistical analyses were performed in R (version 4.0.4).

## Acknowledgements

This work is part of the GENE-SWitCH project that has received funding from the European Union’s Horizon 2020 Research and Innovation Programme under the grant agreement n. 817998. Additional financial support (part of SC salary) was ensured by INRAE, Division of Animal Genetics.

The authors wish to thank Zhan Zhao for help with fecal SCFA measurements; Marie-Stéphane Trottard for help installing software on the Genotoul cluster, Mayrone Mongellaz and Hervé Acloque for help with sample metadata, Jean-Pierre Bidanel and Jordi Estellé for suggestions in the animal experimental design, Michele Hallstead and Joyce Bisdom for help with RNA isolations, Wouter Bakker for help with serum SCFA measurements, Cyril Kurylo for adding functionalities and help running the TAGADA pipeline, and Anne Gabory for useful suggestions for data analyses.

