## Supplementary File 1 for "Differences in maternal diet fiber content influence patterns of gene expression and chromatin accessibility in fetuses and piglets"

### Supplementary Files 1

**Chalabi, Loonen et al. Differences in maternal diet fiber content influence patterns of gene expression and chromatin accessibility in fetuses and piglets.**

#### Supplementary Figures

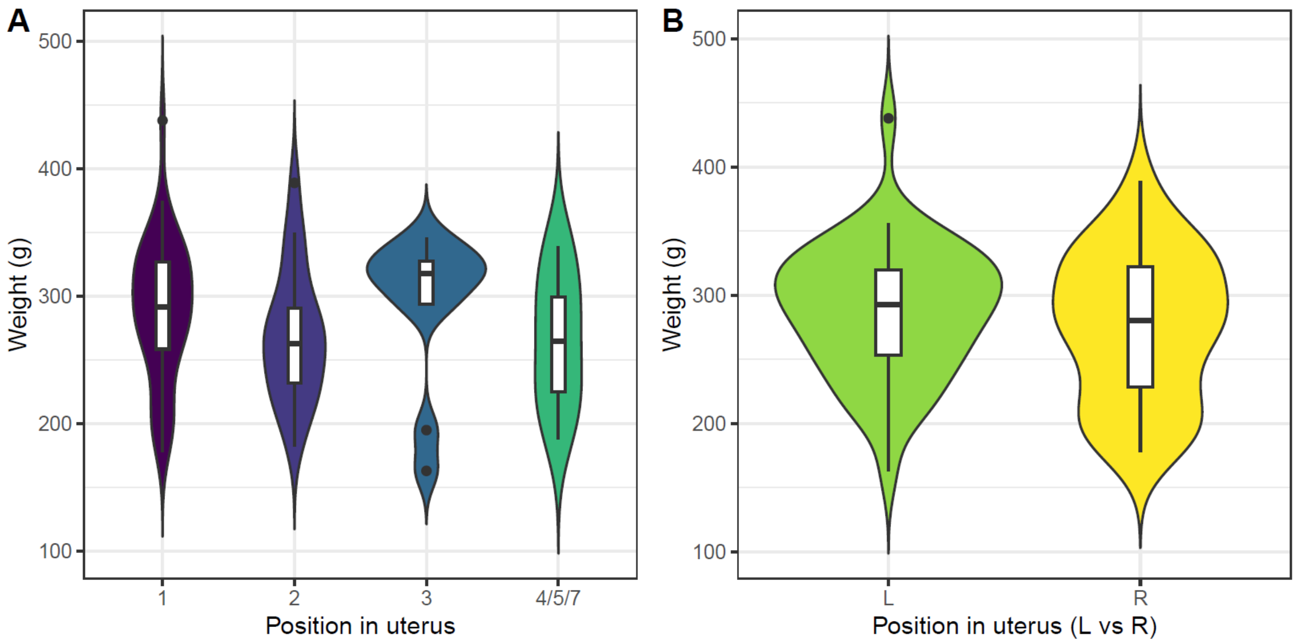


**Supplementary Figure 1**. Impact of fetal intrauterine location on weight according to categorical position counting from the bifurcation of the uterine horns (A), or left versus right uterine horn (B).

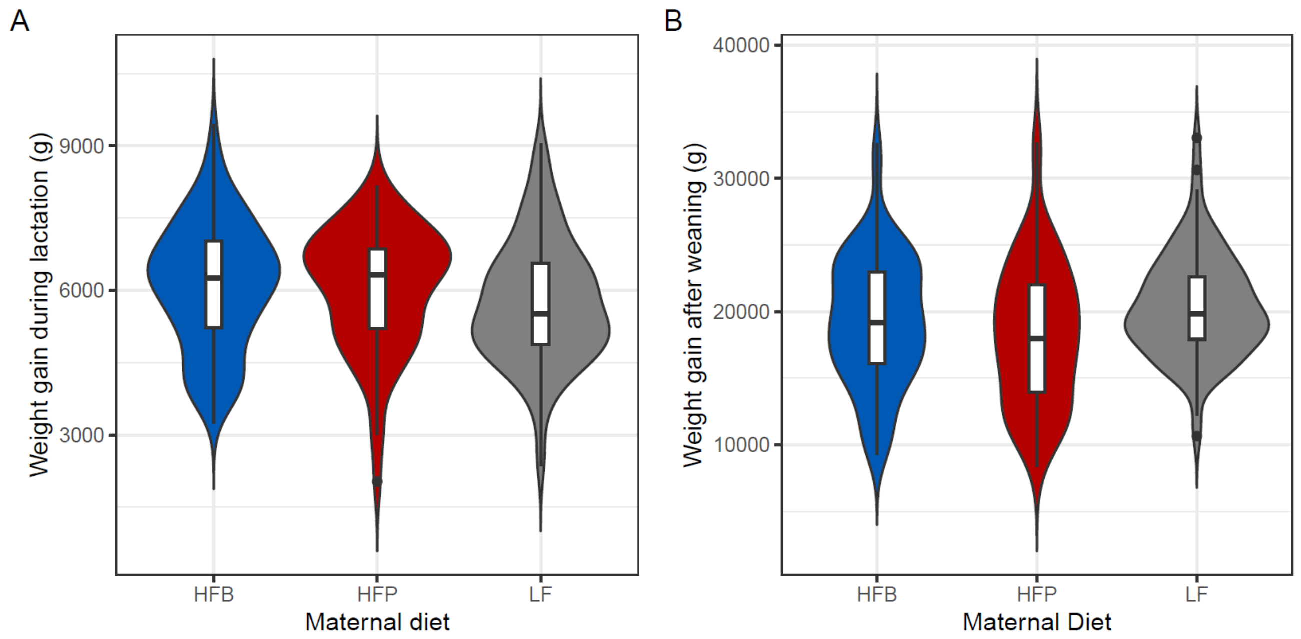


**Supplementary Figure 2**. Violin plots of weight gained (in grams) of study piglets in the 3 maternal diet group (HF-P, red; HF-B, blue; LF; gray) during lactation from birth to weaning (n=284; A), and from weaning to sacrifice (n=248; B). No significant differences were observed between litters.

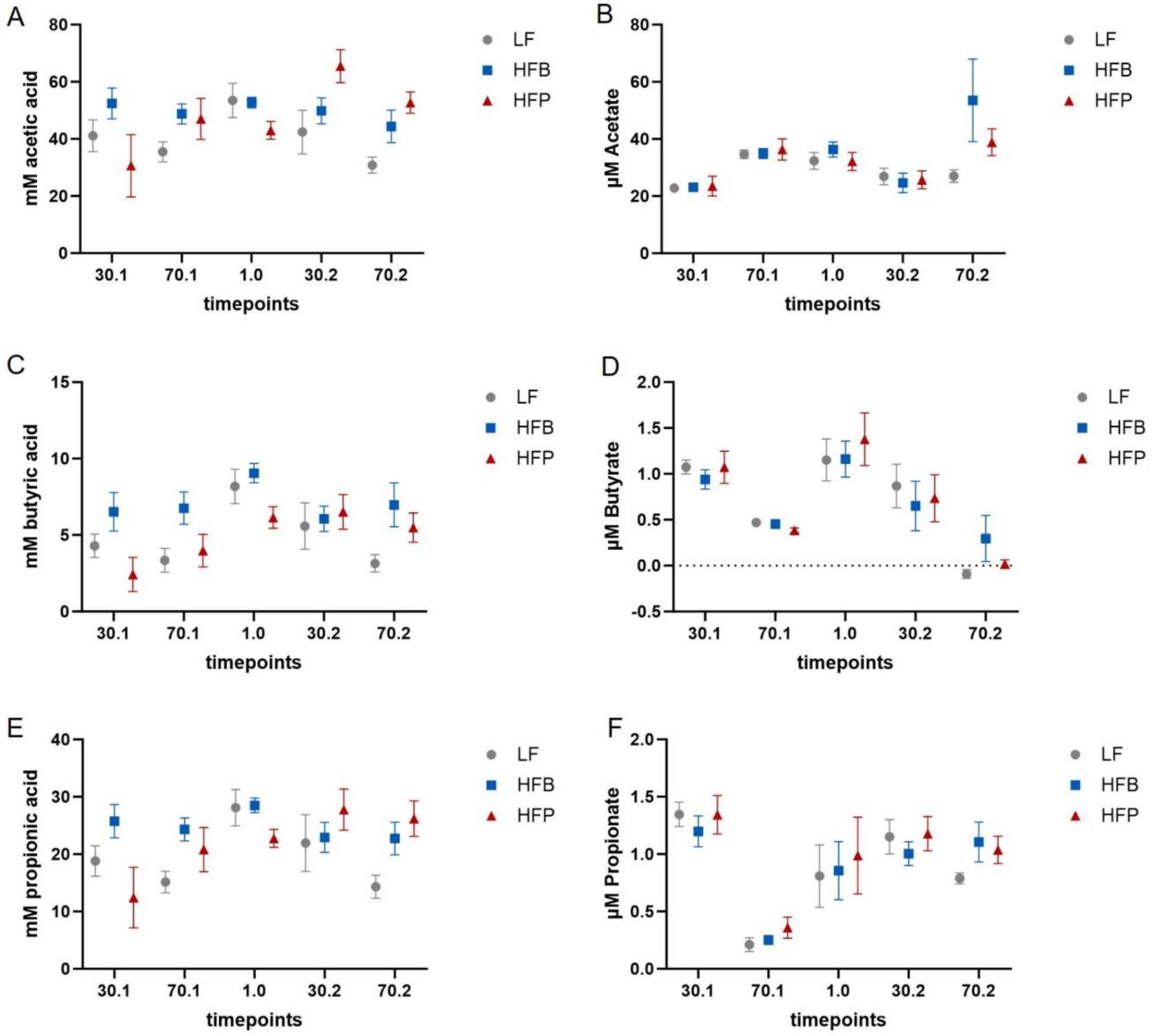


**Supplementary Figure 3**. SCFA levels in sow feces and serum taken on different time points throughout the study. Shown are averages + SEM. Timepoints are day 30 of first pregnancy (30.1), day 70 of first pregnancy (70.1), day 30 of lactation (1.0), day 30 of second pregnancy (30.2) and day 70 of second pregnancy (70.2). Left panels show levels of acidic acid (A), butyric acid (C ) and propionic acid (E) in feces, panels on the right show acetate (B), butyrate (D) and propionate (F) in serum.

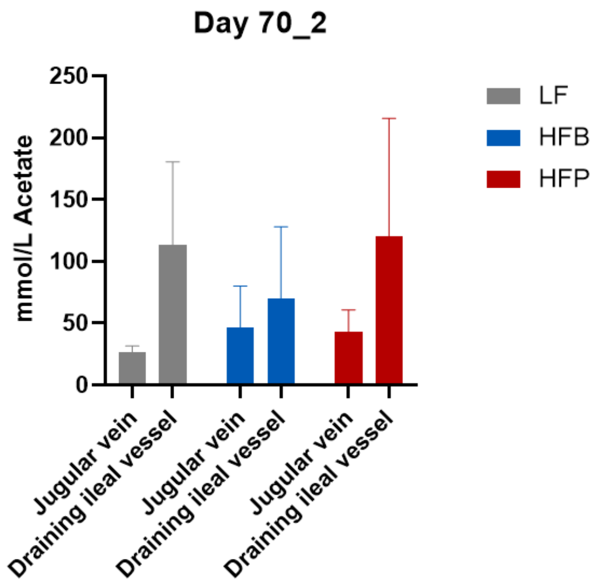


**Supplementary Figure 4**. serum SCFA measures from sows (n=3) per dietary group during second gestation from iliac vein at sacrifice. No statistics were performed due to limited sample size.

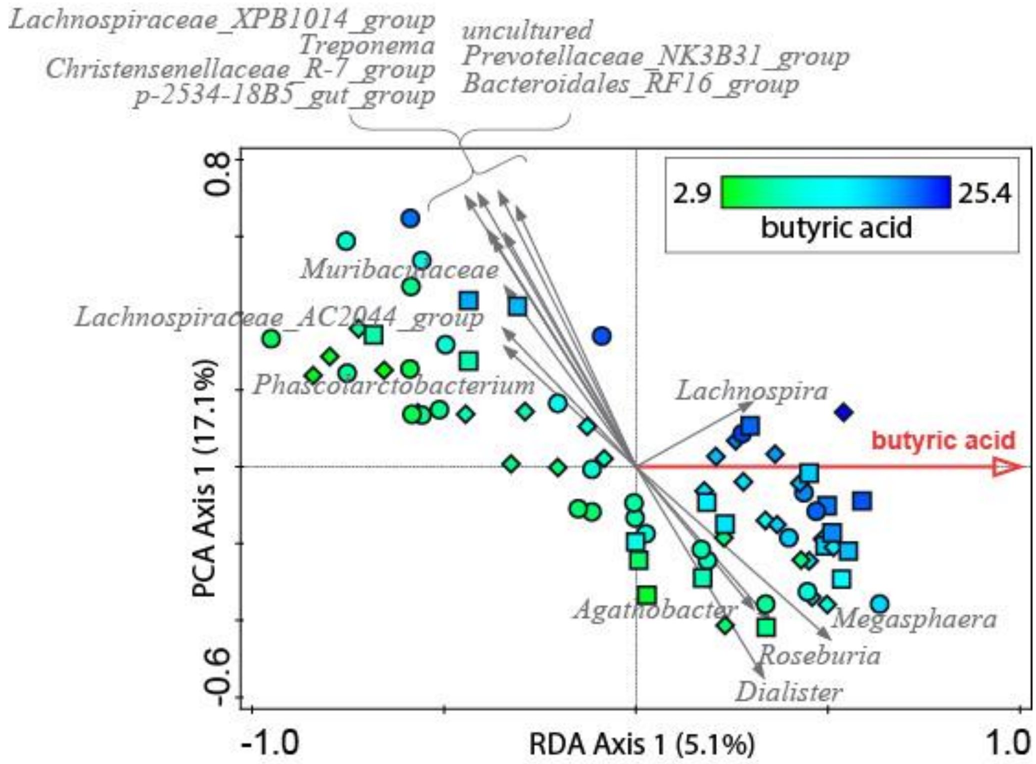


**Supplementary Figure 5**. RDA of piglet microbiota and butyric acids (relative abundances on the genus level as response variables and butyric acids as explanatory variable). Symbols are individual samples, with symbol fill color indicating butyric acid level (color description in the figure legend) and symbol shape indicating diet (squares: HFP, diamonds: HFB, circles: LF). Gray arrows are the 25 best-fitting genera on the horizontal axis, the red arrow indicates increasing butyric acid level in ordination space.

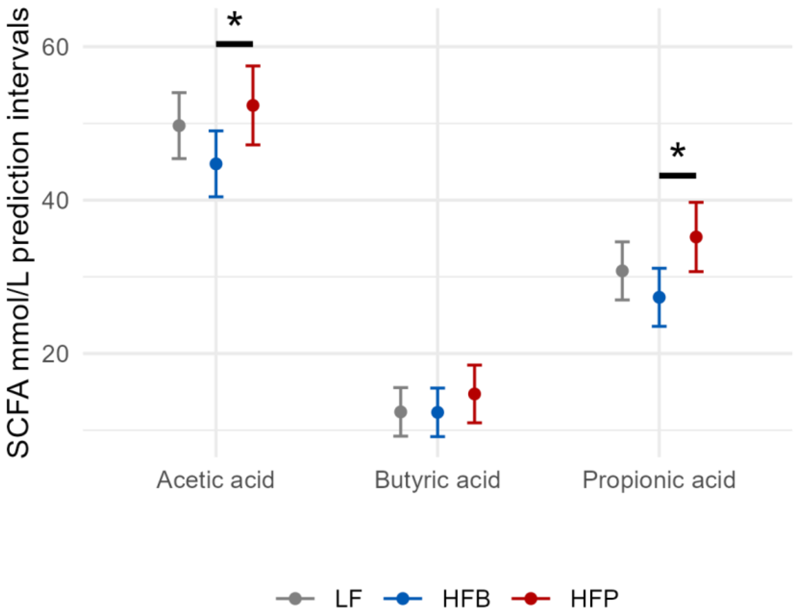


**Supplementary Figure 6**. Linear mixed model adjusted prediction intervals of SCFA measures in piglet feces per maternal dietary group, after accounting for a random litter effect (*n*=4 piglets per litter). Least-squares means were used to test differences between each pair of dietary groups. Significant comparisons ($P < 0.05$) are highlighted with an asterisk (acetic acid HFB vs HFP, P = 0.037; propionic acid HFB vs HFP, P = 0.017).

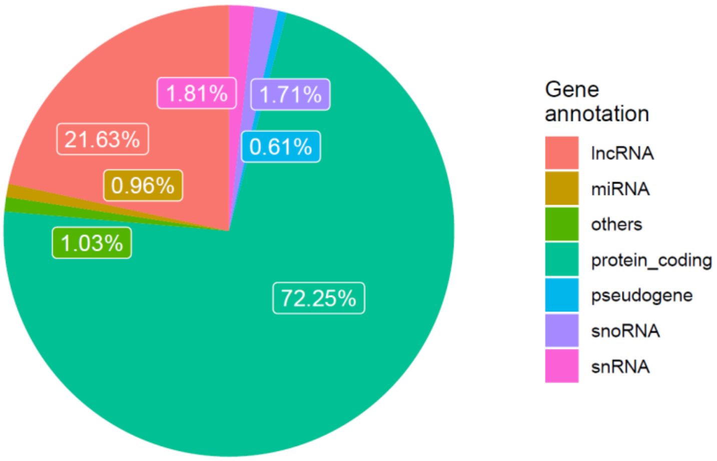


**Supplementary Figure 7**. Biotypes of expressed genes, defined as those with a TPM > 0.1 in at least 2 samples.

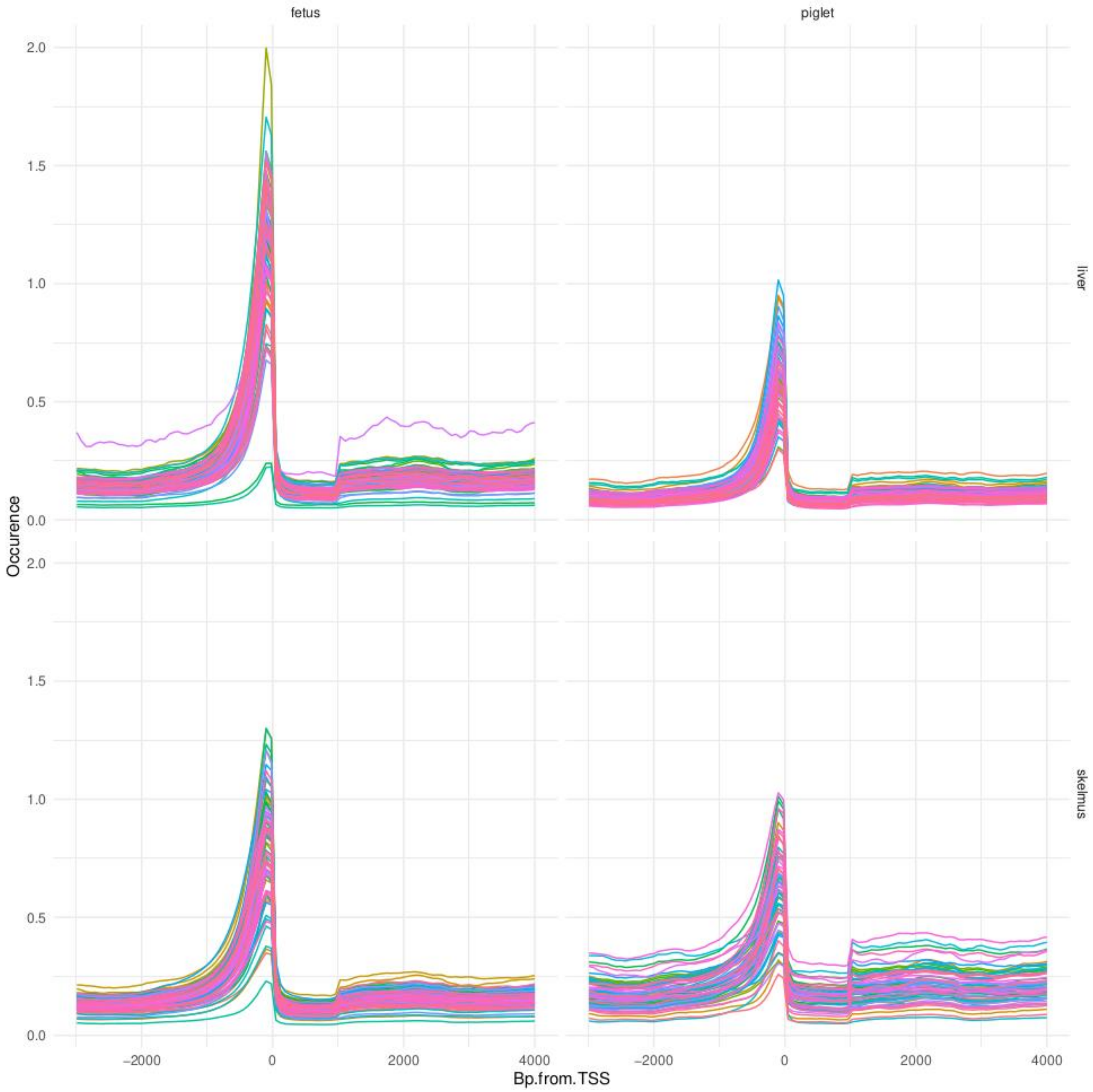


**Supplementary Figure 8**. ATAC-seq mapped read distribution profile within and around annotated genes.

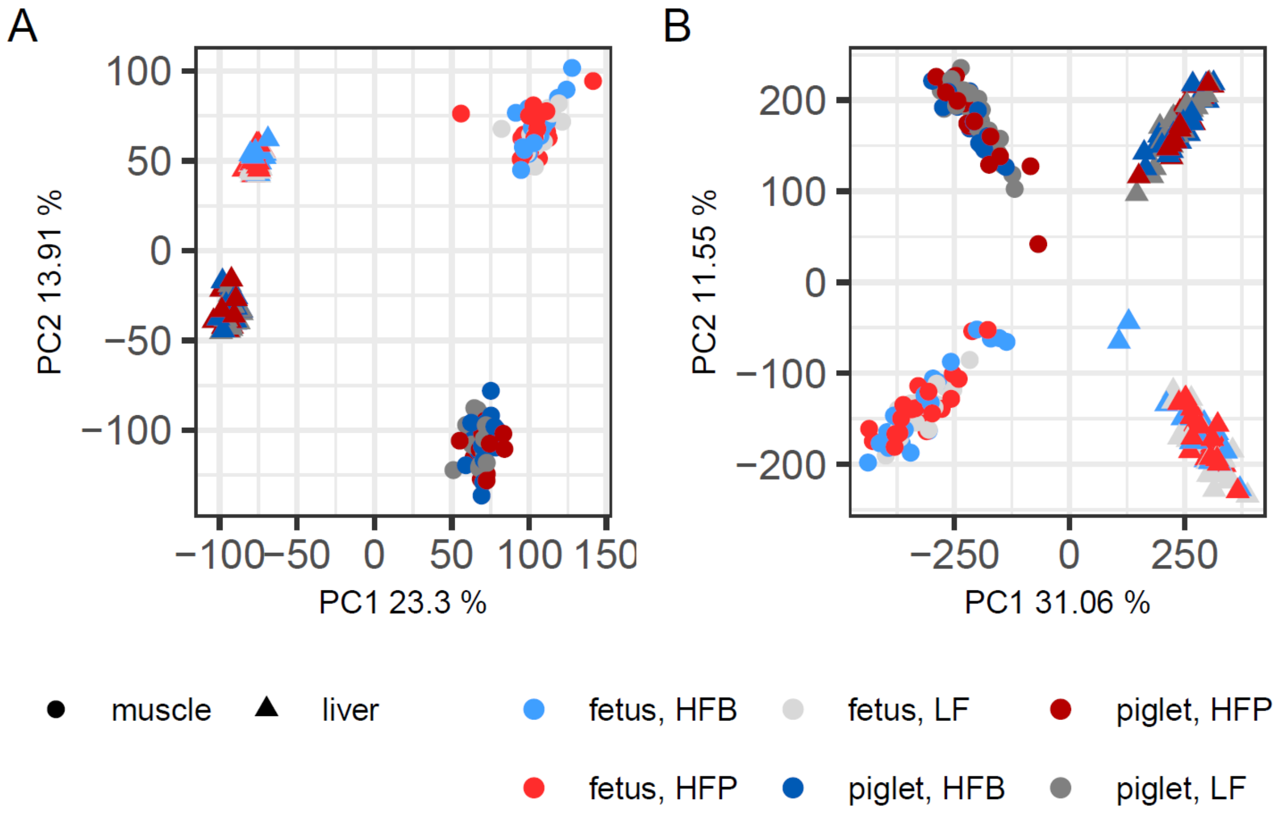


**Supplementary Figure 9**. Principal component analysis (PCA) of normalized ATAC-seq and RNA-seq data. First two axes from a principal components analysis (PCA) for all samples: A) TMM-normalized RNA-seq samples (*n*=296); B) loess-normalized ATAC-seq data (*n*=275). Circles and triangles represent muscle and liver samples, respectively. Colors represent maternal diet groups (blue, HFB; red, HFP; grey, LF), with color saturation reflecting developmental stage (light, fetus; dark, piglet). The percentage of variance explained is indicated next to each PCA axis.

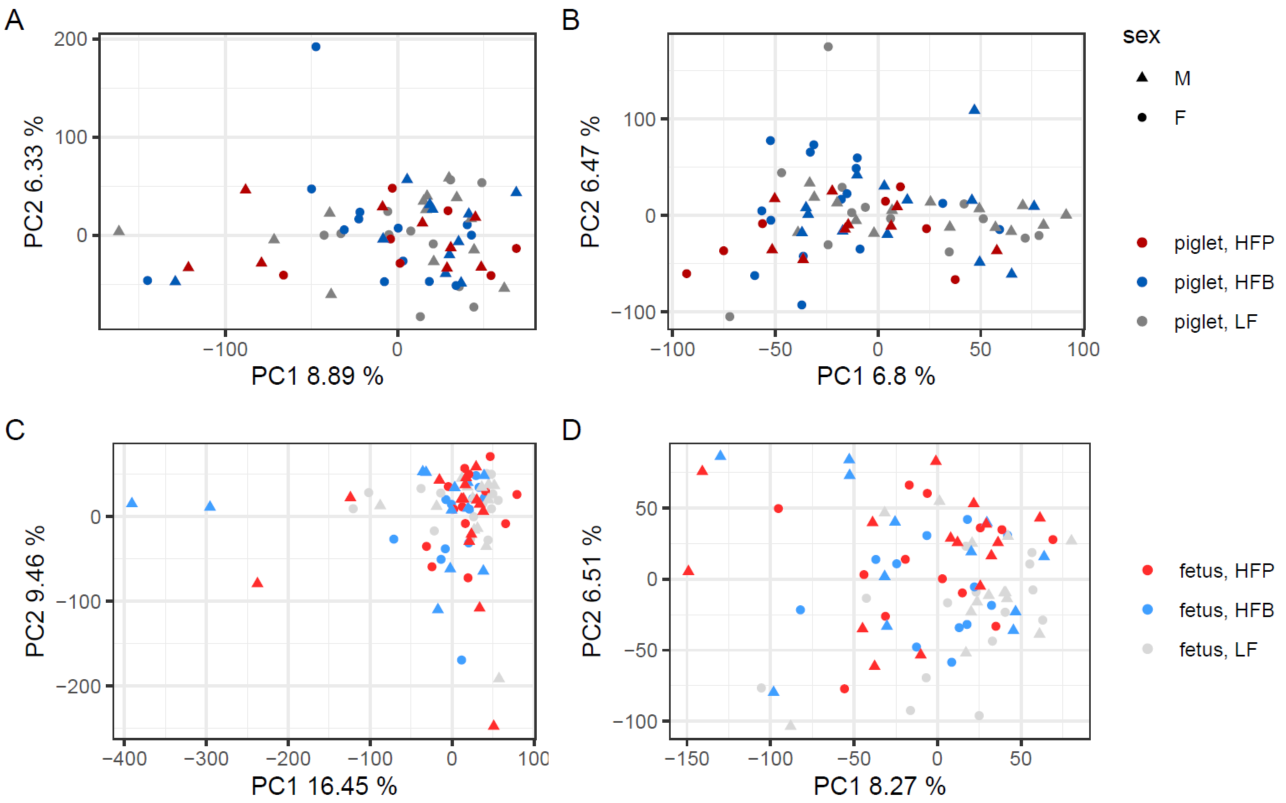


**Supplementary Figure 10**. PCA on TMM-normalized RNA-seq data according to combinations of tissue (liver, muscle), developmental stage (fetus, piglet) and sex, after outlier removal. Colors represent maternal diets and shapes represent sex. (A) piglet muscle samples *n*=64), (B) piglet liver samples (*n*=72), (C) fetus muscle samples (*n*=80), and (D) fetus liver samples (*n*=80 samples). The percentage of variance explained is indicated next to each PCA axis.

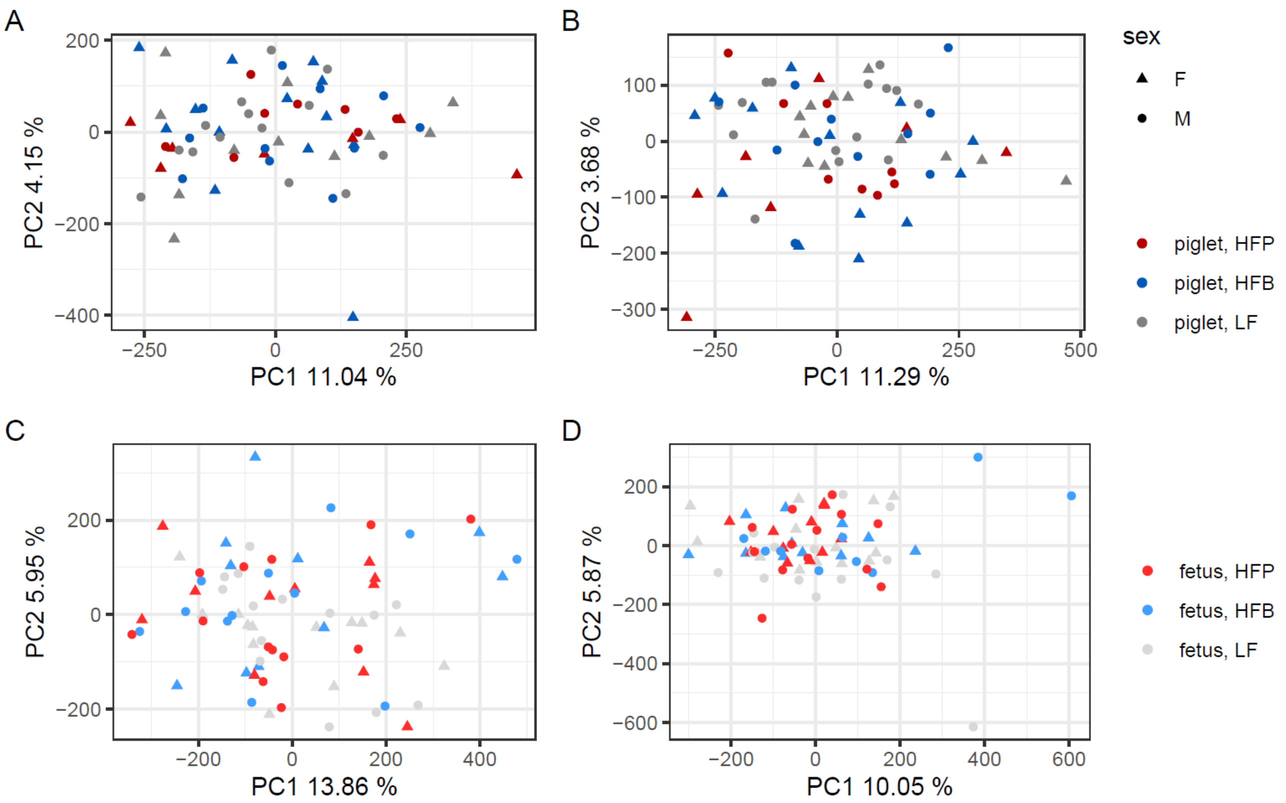


**Supplementary Figure 11**. PCA on loess-normalized ATAC-seq data according to combinations of tissue (liver, muscle), and developmental stage (fetus, piglet) and sex, after outlier removal. Colors represent maternal diets and shapes represent sex. (A) piglet muscle samples (*n*=64), (B) piglet liver samples (*n*=66), (C) fetus muscle samples (*n*=73), and (D) fetus liver samples (*n*=72 samples). The percentage of variance explained is indicated next to each PCA axis.

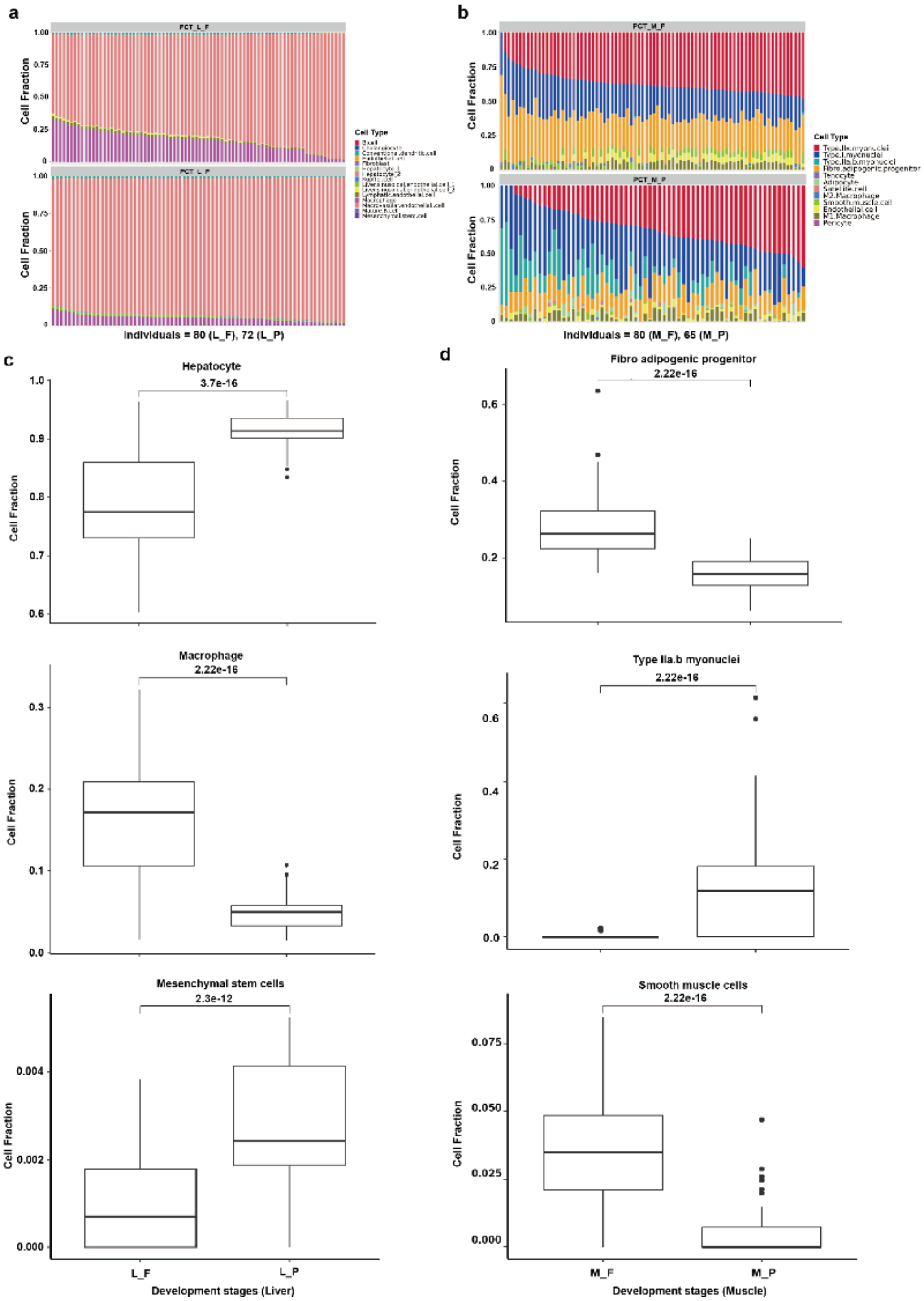


**Supplementary Figure 12. Deconvolution results according to different developmental stages.** (A) Cell component distribution of liver samples in two developmental stages (top: fetal stage, bottom: proliferative stage). (B) Cell component distribution of muscle samples in two developmental stages (top: fetal stage, bottom: proliferative stage). (C) Proportion comparison between two developmental stages within three cell types in liver samples using Student's t-test. (D) Proportion comparison between two developmental stages within three cell types in muscle samples using Student's t-test.

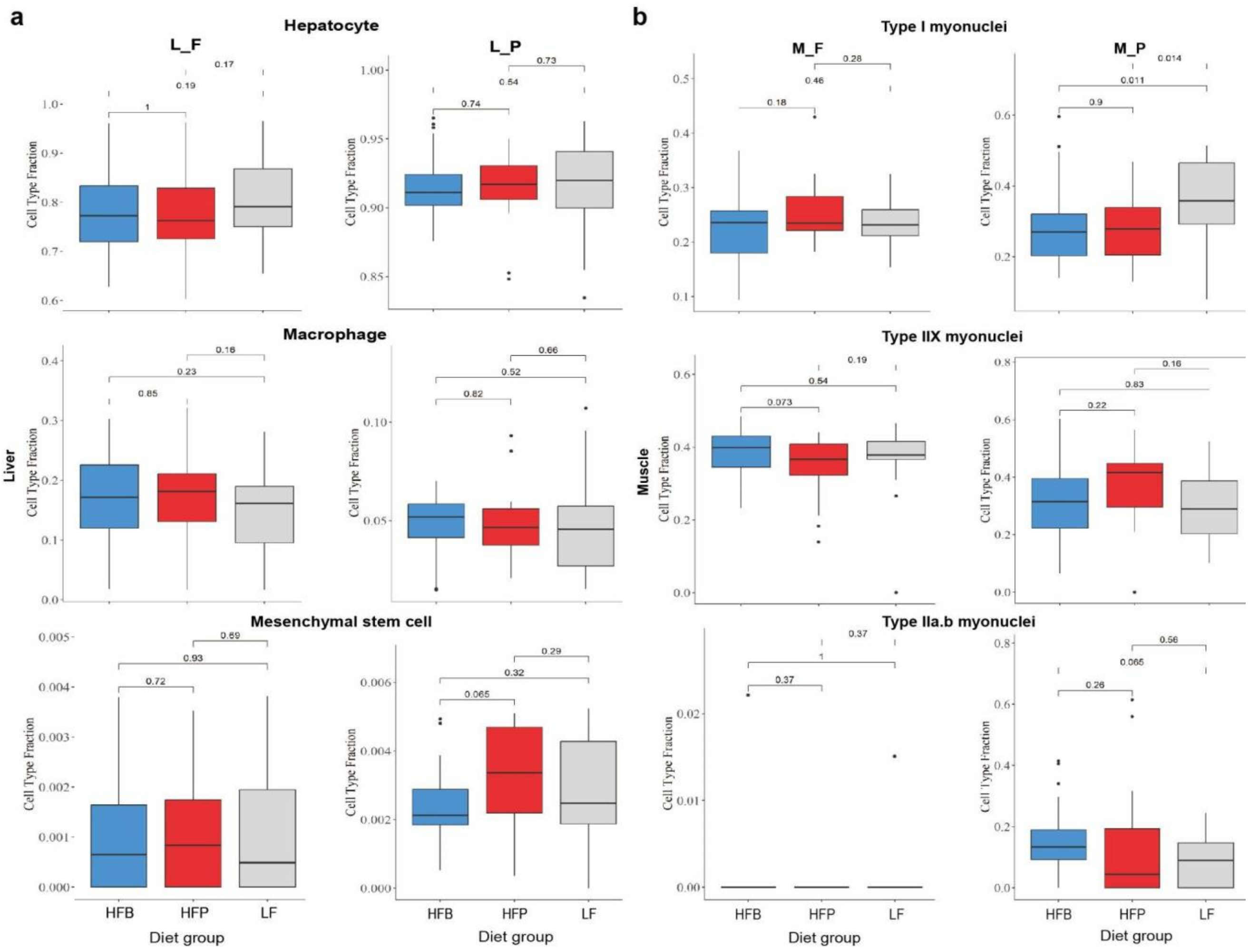


**Supplementary Figure 13. Deconvolution results within important cell types according to different diet groups.** (A) Proportion comparison between three diets within three cell types in liver samples using Student's t-test. (B) Proportion comparison between three diets within three cell types in muscle samples using Student's t-test.

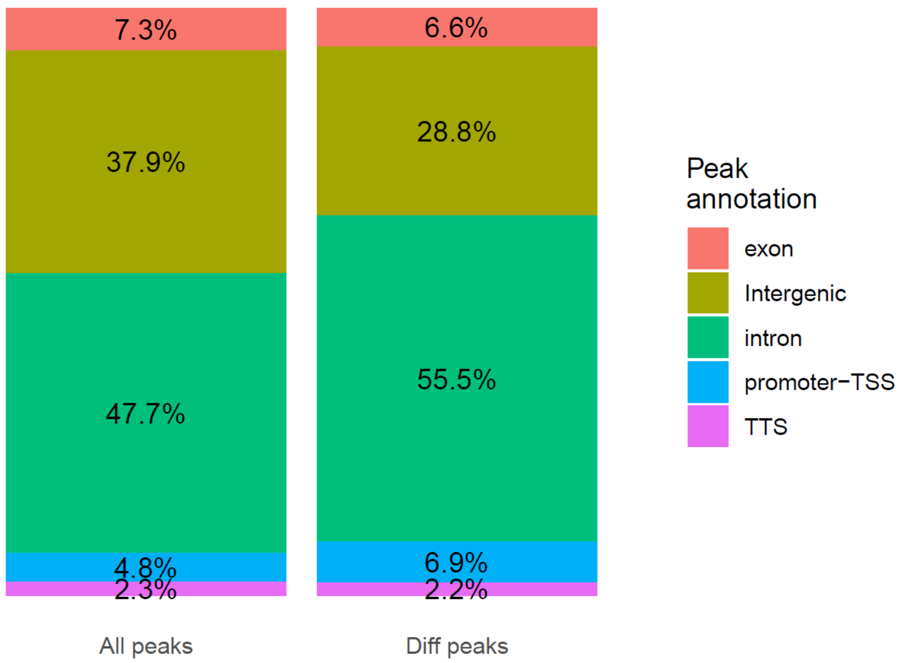


**Supplementary Figure 14**. Classification of accessible chromatin for consensus (left) and differential (right) peaks into genomic annotation categories.

### Supplementary Tables

**Supplementary Table 1**. Composition of experimental diets made by Schothorst feed research (LF = low fiber, HFB = high fiber beet, HFP= high fiber pea). Optiwean was obtained from Trouw Nutrition (the Netherlands); pre-lacto, lacto sow, Wean pellet and Piglet pellet were all obtained from ABZ diervoeding (the Netherlands).

| | LF | HFB | HFP | Pre-lacto | Lacto | Optiwean | Wean pellet | Piglet pellet |
|---|---|---|---|---|---|---|---|---|
| ***Ingredient %*** | | | | | | | | |
| BARLEY cleaned | 44.0 | 26.3 | 26.72 | 14,0 | 10,0 | | 35,0 | 34,0 |
| WHEAT cleaned | 44.0 | 26.3 | 26.72 | 13.24 | 27.95 | | 16.07 | 22.58 |
| CORN | | | | 10.0 | 7.96 | | | 5.5 |
| OAT HULLS | | | | 1.04 | 2.0 | | 3.0 | 1.2 |
| WHEAT bran | | | | 22.39 | 16.0 | | 5.71 | 6.9 |
| Expanded BARLEY | | | | | | | 12.0 | |
| MOLASSES CANE (>47,5% Sug) | 2.2 | 2.5 | 2.5 | 3.0 | 3.0 | | | |
| LINSEED | | | | 1.5 | 1.5 | | 2.0 | |
| TOASTED SOYA BEAN MEAL | | | | | | | 4.42 | |
| HIPRO SOYABEAN MEAL | | | | 2.83 | 8.22 | | | 10.1 |
| SOYA CONCENTRATE | | | | | | | 8.0 | 4.0 |
| POTATO PROTEIN | | | | | | | 1.0 | |
| SUNFLOWER SEED MEAL 38% CP | | | | 2.98 | 3,0 | | | 1.85 |
| SUGAR BEET PULP | | 31.46 | | 5.69 | 2,0 | | 2.0 | 2.0 |
| LIMESTONE | 1.29 | 0.31 | 0.93 | 0.05 | 1.16 | | | |
| BREAD MEAL | | | | 7.46 | 5,0 | | | 5.0 |
| PALMKERNEL FATTY ACIDS | | | | | | | 0.7 | |
| PALM KERNEL MEAL 20% CF | | | | 1.99 | 2.0 | | | |
| RAPESEAD MEAL EXPELLERS | | | | | 1.64 | | | |
| SOYA HULLS 32-36 CF | | | | 4.75 | 2.25 | | | |
| SOYBEAN OIL | 0,0 | 2.09 | 1.59 | 0.79 | 1.0 | | 0.81 | 0.81 |
| SALMON OIL | | | | 0.8 | 0.3 | | 0.4 | 0.3 |
| PALM OIL | | | | 1.0 | 0.5 | | | |
| SOYBEAN MEAL (>48% CP) | 6,0 | 8.62 | 7.5 | | | | | |
| VITAMIN AND MINERAL PREMIXES | 0.4 | 0.4 | 0.4 | | | | | |
| LECITHINE MIX | | | | 1.0 | 0.97 | | | 1,0 |
| WHEY FAT CONCENTRATE | | | | 1.79 | | | 3.89 | |
| CALCIUMFORMIAAT | | | | 0.3 | 0.3 | | 0.1 | 0.2 |
| FORMIC ACID | | | | 0.14 | 0.14 | | | 0.14 |
| LACTIC ACID | | | | 0.49 | 0.49 | | 0.6 | 0.49 |
| BENZOIC ACID | | | | | | | 0.3 | |
| BACT ACID FA | | | | | | | 0.75 | 0.4 |
| PHOSPHORIC ACID | | | | | | | | 0.07 |
| MONOCALCIUM PHOSPHATE | 0.7 | 0.75 | 0.68 | 0.67 | 0.48 | | 0.68 | 0.46 |
| ACID BUF | | | | | | | | 0.47 |
| SALT | 0.15 | 0.1 | | 0.4 | 0.49 | | 0.61 | 0.6 |
| CALCIUMCHLORIDE 78% | | | | 0.1 | | | | |
| SODIUM BICARBONATE | 0.645 | 0.67 | | | 0.14 | | | |
| SODIUM ACETATE | | | 1.0 | | | | | |
| PEA INTERNAL FIBRE | | | 31.46 | | | | | |
| NSP ENZYME ROVABIO EXCEL LC2 | | | | | | | | 0.01 |
| PHYTASE | | | | 0.02 | 0.01 | | | 0.02 |
| METHIONINE (DL,99%) | | | | 0.03 | 0.04 | | 0.16 | 0.15 |
| TRYPTOPHAN (L,98%) | | | | | | | 0.06 | |
| ABZ TRYPTOPHAN (L, 20 %) | | | | | 0.03 | | | 0.15 |
| VALINE (L,99%) | | | | | | | 0.07 | |
| LYSINE HCL (79%) | 0.12 | | | 0.24 | 0.38 | | 0.51 | 0.51 |
| L-THREONINE (98%) | | | | 0.09 | 0.15 | | 0.22 | 0.18 |
| RASPBERRY SWEETENER | | | | | | | 0.25 | 0.13 |
| ABZ WEAN | | | | | | | 0.5 | |
| ABZ VALINE (L,10%) | | | | | | | | 0.38 |
| ABZ sows | 0.5 | 0.5 | 0.5 | 0.5 | 0.5 | | | |
| ABZ PIGLETS e | | | | | | | | 0.4 |
| ABZ lacto | | | | 0.4 | 0.4 | | | |
| M4788 VITAFIX PLUS | | | | 0.1 | | | 0.1 | |
| GLOBAMAX PERFORMANT | | | | | | | 0.1 | |
| CHOLINE-CHLORIDE | | | | 0.07 | | | | |
| ABZ VITAMIN E / VIT E eq | | | | 0.1 | | | | |
| m2606 BIOTIN | | | | 0.05 | | | | |

**Supplementary Table 2.** Chemical composition of experimental diets made by Schothorst feed research (LF = low fiber, HFB = high fiber beet, HFP= high fiber pea). Optiwean was obtained from Trouw Nutrition (the Netherlands); pre-lacto, lacto sow, Wean pellet and Piglet pellet were all obtained from ABZ diervoeding (the Netherlands).

| | LF | HFB | HFP | Pre-lacto | Lacto | Optiwean | Wean pellet | Piglet pellet |
|---|---|---|---|---|---|---|---|---|
| ***Chemical composition*** | | | | | | | | |
| Crude Protein (g/kg) | 131.58 | 130.69 | 120.85 | 130.0 | 153.18 | 165,0 | 161.7 | 167.2 |
| Crude Fat (g/kg) | 24.53 | 39.77 | 37.71 | 67.23 | 55.91 | 60.0 | 55,0 | 40,0 |
| Crude Fibre (g/kg) | 30.46 | 73.77 | 177.19 | 73.5 | 60,0 | 30.0 | 48.22 | 45.0 |
| Ash (g/kg) | 49.83 | 53.4 | 46.96 | 55.99 | 62.21 | 50,0 | 47.57 | 47.37 |
| Dry matter (g/kg) | 875.46 | 881.71 | 894.56 | 889.76 | 888.39 | 900.0 | 898.63 | 893.67 |
| STARCH (g/kg) | 492.22 | 296.27 | 393.71 | 309.05 | 340,0 | 340.0 | 366.33 | 403.45 |
| SUGAR (g/kg) | 37.66 | 56.8 | 44.65 | 71.24 | 53.68 | 60.0 | 61.13 | 38.96 |
| Ca (g/kg) | 7.39 | 6.89 | 6.23 | 5.83 | 9.08 | 6.0 | 4.82 | 5.03 |
| P (g/kg) | 4.65 | 4.11 | 4.91 | 6.04 | 5.56 | 6.0 | 5.26 | 5.15 |
| Digestible Phosphor (g/kg) | | | | 3.7 | 3.35 | 4.0 | 3.85 | 3.55 |
| Sodium (g/kg) | 2.48 | 2.48 | 2.98 | 2.35 | 2.7 | 2.6 | 2.8 | 2.8 |
| K (g/kg) | 5.93 | 6.76 | 8,0 | 8.84 | 8.68 | 8.8 | 7.5 | 7.54 |
| Cl (g/kg) | 2.42 | 1.8 | 1.27 | 5.95 | 5.49 | 5.4 | 6.12 | 5.92 |
| Mg (g/kg) | 1.97 | 2.13 | 1.98 | 2.98 | 2.91 | 1.5 | 1.52 | 1.81 |
| Electrolyte balance (meq) | 191.87 | 230.16 | 298.89 | 165,0 | 190,0 | | 146.97 | 151.3 |
| C18:2 (g/kg) | | | | 22.5 | 21.53 | 19.0 | 19.11 | 19.13 |
| LYS (g/kg) | 5.95 | 6.06 | 5.82 | 6.86 | 9.22 | 12.3 | 11.45 | 11.37 |
| MET (g/kg) | 2.01 | 1.99 | 1.71 | 2.29 | 2.7 | 4.54 | 4.0 | 3.9 |
| Cu (mg/kg) | 19.62 | 20.76 | 20.76 | 21.53 | 21.71 | | 146.59 | 96.4 |
| Fe (mg/kg) | 308.59 | 438.74 | 314.78 | 345.07 | 321.68 | 125,0 | 182.85 | 228.66 |
| Mn (mg/kg) | 66.05 | 74.3 | 61.95 | 86.58 | 83.49 | 50.0 | 72.05 | 57.36 |
| Zn (mg/kg) | 131.18 | 126.37 | 130.69 | 140.86 | 137.84 | 105.0 | 130.29 | 131.31 |
| Cu (mg/kg) | | | | 5.03 | 5.03 | 150.0 | 140.7 | 90.45 |
| Se (mg/kg) | | | | 0.3 | 0.3 | 0.4 | 0.4 | 0.3 |
| Mn (mg/kg) | | | | 20.1 | 20.1 | | | |
| Zn, organic (mg/kg) | | | | 50.25 | 50.25 | | | |
| Se, Selenomethionine (mg/kg) | | | | 0.2 | 0.2 | | 0.15 | 0.15 |
| Cu, organic (mg/kg) | | | | 10.05 | 10.05 | | | |
| Potassium iodide (mg/kg) | | | | | | 2.5 | | |
| Betainehydrochloride (mg/kg) | | | | | | 150,0 | | |
| Lactose (g/kg) | | | | | | 75.0 | | |
| Vit, A (ie) | | | | 12060.3 | 12060.3 | 15000.0 | 15075.38 | 15075.38 |
| Vit, D3 (ie) | | | | 1005.03 | 1005.03 | 2000.0 | 1005.03 | 1005.03 |
| Vit, D3 + D3OH (ie) | | | | 2010.05 | 2010.05 | | 2010.05 | 2010.05 |
| Vit, E + EQ (mg/kg) | | | | 120,0 | 90.45 | 200.0 | 160.8 | 100.5 |
| Vit, B1, Thiamin (mg/kg) | | | | 1.01 | 1.01 | 2.0 | 1.01 | 1.01 |
| Vit, B2 (mg/kg) | | | | 6.03 | 6.03 | 6.0 | 5.03 | 5.03 |
| Vit, B5, Pantothenic acid (mg/kg) | | | | 16.42 | 16.42 | 15.0 | 10.92 | 10.94 |
| Vit, B3, Niacin (mg/kg) | | | | 25.13 | 25.13 | 50.0 | 35.18 | 35.18 |
| Vit, B6 (mg/kg) | | | | 1.01 | 1.01 | 3.0 | 1.01 | 1.01 |
| Vit, B9/B11, Folic acid (mg/kg) | | | | 3.02 | 3.02 | 1.0 | 0.5 | 0.4 |
| Vit, B12 (mg/kg) | | | | 0.03 | 0.03 | 40,0 | 0.03 | 0.03 |
| Vit, K3 (mg/kg) | | | | 2.01 | 2.01 | 2.0 | 2.51 | 2.51 |
| Vit, B4, Choline chloride (mg/kg) | | | | 950.0 | 603.02 | | 201.01 | 100.5 |
| Vit, C (mg/kg) | | | | 50.25 | 50.25 | 50,0 | | |
| Vit, H, Biotin (mg/kg) | | | | 0.4 | 0.3 | 150.0 | | |
| Net Energy gestation sows (x100) | | | | 111.36 | 110.39 | | | |
| Net energy lactating sows (x100) | | | | 110.0 | 109.0 | | | |
| Net Energy piglets (x100) | | | | | | | 108.0 | 107.15 |
| fCHO-s (g/kg) | 88.34 | 238.84 | 163.94 | | | | | |
| iCHO-s (g/kg) | 55.96 | 69.6 | 87.56 | | | | | |
| Moisture (g/kg) | 124.54 | 118.29 | 105.44 | | | | | |
| S inorg (g/kg) | 0.32 | 0.76 | 0.72 | | | | | |
| S organic (g/kg) | 1.12 | 1.02 | 0.94 | | | | | |
| S total (g/kg) | 1.53 | 1.88 | 1.75 | | | | | |
| dEBs (meq) | 94.41 | 110.88 | 187.78 | | | | | |
| ABC4 (meq) | 423.89 | 259.49 | 297.12 | | | | | |
| CYS (g/kg) | 2.61 | 2.29 | 2.16 | | | | | |
| M+C (g/kg) | 4.62 | 4.28 | 3.87 | | | 7.64 | | |
| THR (g/kg) | 4.29 | 4.92 | 4.17 | | | 8.66 | | |
| TRP (g/kg) | 1.63 | 1.58 | 1.44 | | | 2.77 | | |
| VAL (g/kg) | 6.01 | 6.36 | 5.61 | | | | | |
| ILE (g/kg) | 4.79 | 5.0 | 4.68 | | | | | |
| LEU (g/kg) | 9.03 | 8.99 | 8.52 | | | | | |
| HIS (g/kg) | 3.13 | 3.42 | 2.96 | | | | | |
| SID Lys (g/kg) | 5.14 | 4.74 | 4.92 | 5.58 | 7.9 | 11.0 | 10.05 | 10.07 |
| SID Met (g/kg) | 1.76 | 1.63 | 1.48 | 1.91 | 2.32 | 3.63 | 3.61 | 3.61 |
| SID Met+Cys (g/kg) | 4,0 | 3.45 | 3.28 | 3.69 | 4.43 | 6.38 | 5.82 | 5.82 |
| SID Thr (g/kg) | 3.66 | 3.42 | 3.5 | 3.69 | 5.14 | 6.93 | 6.33 | 6.33 |
| SID Trp (g/kg) | 1.39 | 1.24 | 1.2 | 1.14 | 1.5 | 2.2 | 2.11 | 2.11 |
| SID Valine (g/kg) | 5.18 | 4.82 | 4.77 | 4.36 | 5.36 | | 6.52 | 6.52 |
| SID LYS-s/E-gestate (ratio) | 0.52 | 0.52 | 0.48 | | | | | |
| SID ILE-s (g/kg) | 4.19 | 4.05 | 4.04 | | | | | |
| SID LEU-s (g/kg) | 7.83 | 7.23 | 7.28 | | | | | |
| SID HIS-s (g/kg) | 2.76 | 2.73 | 2.59 | | | | | |
| Net Energy gest sows (MJ/kg) | 9.88 | 9.14 | 10.25 | | | | | |
| SID MET/SID LYS-s (ratio) | 0.34 | 0.34 | 0.3 | | | | | |
| SID M+C/SID LYS-s (ratio) | 0.78 | 0.73 | 0.67 | | | | | |
| SID THR/SID LYS-s (ratio) | 0.71 | 0.72 | 0.71 | | | | | |
| SID TRP/SID LYS-s (ratio) | 0.27 | 0.26 | 0.24 | | | | | |
| SID VAL/SID LYS-s (ratio) | 1.01 | 1.02 | 0.97 | | | | | |
| SID ILE/SID LYS-s (ratio) | 0.82 | 0.85 | 0.82 | | | | | |
| SID LEU/SID LYS-s (ratio) | 1.52 | 1.53 | 1.48 | | | | | |
| SID HIS/SID LYS-s (ratio) | 0.54 | 0.58 | 0.53 | | | | | |
| NSP-s total (g/kg) | 135.01 | 300.14 | 240.66 | | | | | |
| 6-Fytase EC 3,1,3,26 (U/kg) | | | | | | 1200,0 | | |
| Endo-1,3(4)-bèta-glucanase EC 3,2,1,6 (U/kg) | | | | | | 152.0 | | |
| Endo-1,4-bèta-xylanase EC 3,2,1,8 (U/kg) | | | | | | 1220.0 | | |
| B. licheniformis + B. subtilis (1:1) (10^6 CFU/kg) | | | | | | 1280.0 | | |
| Butylhydroxytoluene Antioxidant (mg/kg) | | | | | | 150.0 | | |
| Propylgallate (mg/kg) | | | | | | 8.64 | | |
| Na+K-Cl (mEq/kg) | | | | | | 186.0 | | |

**Supplementary Table 3**. Results of fetus weight analyses, based on ANOVA Type III test results and a linear model with a random litter effect. *In utero* position was coded as a categorical variable. Analysis of Deviance tables include the $\chi^2$ test statistic, corresponding degrees of freedom (df), and *P*-value. Significance codes are as follows: "***": $P < 0.0001$, "**": $P < 0.001$, "**": $P < 0.01$, "*" : $P < 0.05$, ".": $P < 0.1$. Results corresponding to the effect of maternal diet are highlighted in boldface.

| | | $\chi^2$ | df | Pr(>Chi$^2$) |
|---|---|---|---|---|
| Weight, fetuses (*n*=80) | (Intercept) | 113.2928 | 1 | < 2e-16 *** |
| | Sex | 3.5167 | 1 | 0.06075 |
| | Maternal breed | 0.7635 | 1 | 0.38224 |
| | Paternal breed | 3.3985 | 1 | 0.06526 . |
| | Position *in utero* | 5.3876 | 5 | 0.37043 |
| | **Maternal diet** | **0.7172** | **2** | **0.69866** |

**Supplementary Table 4**. Results of piglet weight analyses, based on ANOVA Type III test results; all linear models include a random litter effect. Analysis of Deviance tables include the $\chi^2$ test statistic, corresponding degrees of freedom (df), and *P*-value. Significance codes are as follows: "***": $P < 0.0001$, "**": $P < 0.001$, "**": $P < 0.01$, "*" : $P < 0.05$, ".": $P < 0.1$. Results corresponding to the effect of maternal diet are highlighted in boldface.

| | | $\chi^2$ | df | Pr(>Chi$^2$) |
|---|---|---|---|---|
| Birth weight, all piglets born (*n*=392) | (Intercept) | 27.7014 | 1 | 1.416e-07 *** |
| | Round | 1.1372 | 1 | 0.28624 |
| | Maternal breed | 0.2665 | 1 | 0.60566 |
| | Parity | 0.3833 | 1 | 0.53586 |
| | Litter size | 5.2794 | 1 | 0.02158 * |
| | **Maternal diet** | **4.1119** | **2** | **0.12797** |
| Birth weight, post-weaned piglets fed by birth mother (*n*=253) | (Intercept) | 16.6684 | 1 | 4.452e-05 *** |
| | Round | 0.0140 | 1 | 0.90567 |
| | Maternal breed | 0.0627 | 1 | 0.80223 |
| | Parity | 0.1446 | 1 | 0.70380 |
| | Litter size | 4.1503 | 1 | 0.04163 * |
| | **Maternal diet** | **2.0156** | **2** | **0.36502** |
| Weaning weight, post-weaned piglets fed by birth mother (*n*=253) | (Intercept) | 0.3472 | 1 | 0.55572 |
| | Round | 0.7321 | 1 | 0.39222 |
| | Maternal breed | 0.4736 | 1 | 0.49134 |
| | Parity | 6.3456 | 1 | 0.01177 * |
| | Litter size | 6.0146 | 1 | 0.01419 * |
| | Age at weaning | 3.3067 | 1 | 0.06900 . |
| | Number of weaned piglets | 0.1166 | 1 | 0.73271 |
| | **Maternal diet** | **1.2414** | **2** | **0.53756** |
| Weight gain from birth to weaning, post-weaned piglets fed by birth mother (*n*=253) | (Intercept) | 0.0768 | 1 | 0.781638 |
| | Round | 2.1236 | 1 | 0.145046 |
| | Maternal breed | 1.3988 | 1 | 0.236928 |
| | Parity | 10.0894 | 1 | 0.001491 ** |
| | Litter size | 5.0458 | 1 | 0.024686 * |
| | Age at weaning | 5.0652 | 1 | 0.024411 * |
| | Number of weaned piglets | 0.0013 | 1 | 0.971363 |
| | **Maternal diet** | **4.7586** | **2** | **0.092614 .** |

**Supplementary Table 5**. Number of samples for each combination of tissue, stage, and maternal diet used for RNA-seq and ATAC-seq assays. The total number of samples across all combinations is n=296 and n=275 for RNA-seq and ATAC-seq, respectively. In total, n=8 RNA-seq samples were removed due to an atypically low average number of reads, and n=29 ATAC-seq samples were removed as outliers that failed quality control checks: weak TSS enrichment (n=4), FrIP score <10% (n=8), failing GC content (n=3), high percentage of base-pair trimming (n=14), and high duplication (n=3).

| | | LF | | HFP | | HFB | |
|---|---|---|---|---|---|---|---|
| | | fetus | piglet | fetus | piglet | fetus | piglet |
| RNA-seq | muscle | 28 | 24 | 28 | 16 | 24 | 24 |
| | liver | 28 | 28 | 28 | 16 | 24 | 28 |
| ATAC-seq | muscle | 27 | 25 | 24 | 15 | 22 | 24 |
| | liver | 28 | 27 | 24 | 15 | 20 | 24 |

**Supplementary Table 6**. Number of significantly (FDR < 5%) differentially expressed genes in each combination of developmental stage (fetus, piglet) and tissue (liver, muscle), after accounting for sex, maternal breed, paternal breed, and a random litter effect. Results are shown for each pairwise diet comparison (HF-P vs LF, HF-B vs LF, and HF-B vs HF-B), as well as for the remaining fixed effects in each model. No paternal breed effect could be estimated for piglets, as all piglet fathers were of a single breed.)

| | | | **Diet** | | | **Sex** | **Maternal breed** | **Paternal breed** |
|---|---|---|---|---|---|---|---|---|
| | | | **HFP *vs* LF** | **HFB *vs* LF** | **HFB *vs* HFP** | **M *vs* F** | **70 *vs* 60** | **Y *vs* T** |
| **Fetus** | **Liver** | **Up** | 0 | 0 | 0 | 44 | 236 | 255 |
| | | **Down** | 0 | 0 | 0 | 74 | 130 | 357 |
| | **Muscle** | **Up** | 0 | 0 | 0 | 20 | 1 | 168 |
| | | **Down** | 0 | 0 | 0 | 21 | 1 | 217 |
| **Piglets** | **Liver** | **Up** | 0 | 0 | 0 | 162 | 10 | NA |
| | | **Down** | 0 | 0 | 0 | 146 | 6 | |
| | **Muscle** | **Up** | 0 | 0 | 0 | 23 | 19 | |
| | | **Down** | 0 | 0 | 0 | 28 | 5 | |

**Supplementary Table 7.** Number of significantly (FDR < 5%) differentially accessible peaks in each combination of developmental stage (fetus, piglet) and tissue (liver, muscle), after accounting for sex, maternal breed, paternal breed, and a random litter effect. Results are shown for each pairwise diet comparison (HFP vs LF, HFB vs LF, and HFB vs HFB), as well as for the remaining fixed effects in each model. No paternal breed effect could be estimated for piglets, as all piglet fathers were of a single breed.

| | | | **Diet** | | | **Sex** | **Maternal breed** | **Paternal breed** | |
|---|---|---|---|---|---|---|---|---|---|
| | | | **HFP *vs* LF** | **HFB *vs* LF** | **HFB *vs* HFP** | **M *vs* F** | **70 *vs* 60** | **Y *vs* T** | **Total** |
| **Fetus** | **Liver** | **Open** | 0 | 1 | 1 | 279 | 18 | 2,317 | 197,695 |
| | | **Closed** | 0 | 0 | 2 | 1,425 | 7 | 492 | |
| | **Muscle** | **Open** | 1 | 3 | 1 | 158 | 79 | 362 | 200,385 |
| | | **Closed** | 1 | 1 | 1 | 397 | 55 | 805 | |
| **Piglets** | **Liver** | **Open** | 6 | 1 | 3 | 413 | 19,151 | NA | 196,375 |
| | | **Closed** | 5 | 7 | 11 | 2,141 | 1,894 | | |
| | **Muscle** | **Open** | 0 | 527 | 7 | 191 | 2,694 | | 173,099 |
| | | **Closed** | 2 | 119 | 6 | 517 | 6,472 | | |
| **Total** | | | 15 | 659 | 32 | 5,521 | 30,370 | | |
